# Collagen (I) homotrimer potentiates the osteogenesis imperfecta (oim) mutant allele and reduces survival in male mice

**DOI:** 10.1101/2020.07.13.198283

**Authors:** Katie J. Lee, Lisa Rambault, George Bou-Gharios, Peter D. Clegg, Riaz Akhtar, Gabriela Czanner, Rob van ‘t Hof, Elizabeth G. Canty-Laird

## Abstract

Type I collagen is the major structural component of bone where it exists as an (α1)_2_(α2)_1_ heterotrimer in all vertebrates. The osteogenesis imperfecta (oim) mouse model comprising solely homotrimeric (α1)_3_ type I collagen, due to a dysfunctional α2 chain, has a brittle bone phenotype implying that the heterotrimeric form is required for physiological bone function. However, humans with rare null alleles preventing synthesis of the α2 chain have connective tissue and cardiovascular abnormalities (cardiac valvular Ehlers Danlos Syndrome), without evident bone fragility. Conversely a prevalent human single nucleotide polymorphism leading to increased homotrimer synthesis is associated with osteoporosis. Whilst the oim line is well-studied, whether homotrimeric type I collagen is functionally equivalent to the heterotrimeric form in bone has not been demonstrated. Col1a2 null and oim mouse lines were used in this study and bones analysed by microCT and 3-point bending. RNA was also extracted from heterozygote tissues and allelic discrimination analyses performed using qRT-PCR. Here we comprehensively show for the first time that mice lacking the α2(I) chain do not have impaired bone biomechanical or structural properties, unlike oim homozygous mice. However Mendelian inheritance was affected in male mice of both lines and male mice null for the α2 chain exhibited age-related loss of condition. The brittle bone phenotype of oim homozygotes could result from detrimental effects of the oim mutant allele, however, the phenotype of oim heterozygotes is known to be less severe. We used allelic discrimination to show that the oim mutant allele is not downregulated in heterozygotes. We then tested whether gene dosage was responsible for the less severe phenotype of oim heterozygotes by generating compound heterozygotes. Data showed that compound heterozygotes had impaired bone structural properties as compared to oim heterozygotes, albeit to a lesser extent than oim homozygotes. Hence, we concluded that the presence of heterotrimeric collagen-1 in oim heterozygotes alleviates the effect of the oim mutant allele but a genetic interaction between homotrimeric collagen-1 and the oim mutant allele leads to bone fragility.

## Introduction

Type I collagen is the major structural component of vertebrate tissues, where it exists as insoluble fibrils formed from arrays of trimeric collagen molecules. In tetrapods, type I collagen molecules are predominantly (α1)_2_(α2)_1_ heterotrimers derived from the polypeptide gene products of the Col1a1 and Col1a2 genes. Trimeric type I procollagen molecules contain a central triple-helical domain flanked by globular N- and C-propeptide regions, that are proteolytically removed to facilitate fibrillogenesis (1). Procollagen molecules are synthesised in the endoplasmic reticulum of the secretory pathway where individual molecules first associate via the C-propeptides. C-propeptide association facilitates chain registration and folding of the triple helical domain into a right-handed triple helix. The type-specific assembly of fibrillar procollagens has been attributed to defined amino acid sequences within the C-propeptide including a chain recognition sequence (2), specific stabilising residues (3) and a cysteine code (4), although none fully explain the preferential heterotrimerisation of type I procollagen.

Abnormal type I collagen (α1)_3_ homotrimer, derived from COL1A1 alone, is genetically or biochemically associated with common age-related human diseases including osteoporosis (5), osteoarthritis (6–8), intervertebral disc degeneration (9), arterial stiffening (10), cancer (11), liver fibrosis (12) and Dupuytren’s contracture (13). The homotrimeric form is resistant to mammalian collagenases (14) and alterations in collagen crosslinking have been reported in the osteogenesis imperfecta murine (oim) model, which lacks a functional α2(I) chain (15–17). Hence the presence of homotrimeric collagen (I) in human disease may alter the ability of tissues to respond to changing physiological demands by slowing remodelling and altering tissue mechanics.

The oim mutation is a deletion of a single guanidine residue, causing a frameshift that alters the last 47 amino acids, and reportedly adds an additional residue to the α2 chain of type I procollagen (Fig. 1A). The mutant α2 chain cannot be incorporated into trimers hence giving rise to solely homotrimeric collagen-1 in oim homozygotes. Homozygous oim/oim mice have osteopenia, progressive skeletal deformities, spontaneous fractures, cortical thinning and small body size (18) with corresponding alterations to bone structure and material properties (19, 20). Homozygous oim mice also have a reduced circumferential breaking strength and greater compliance of aortae (16, 21), reduced ultimate stress and strain tendon for tendon (22) and kidney glomerulopathy (23).

**Figure 1.**
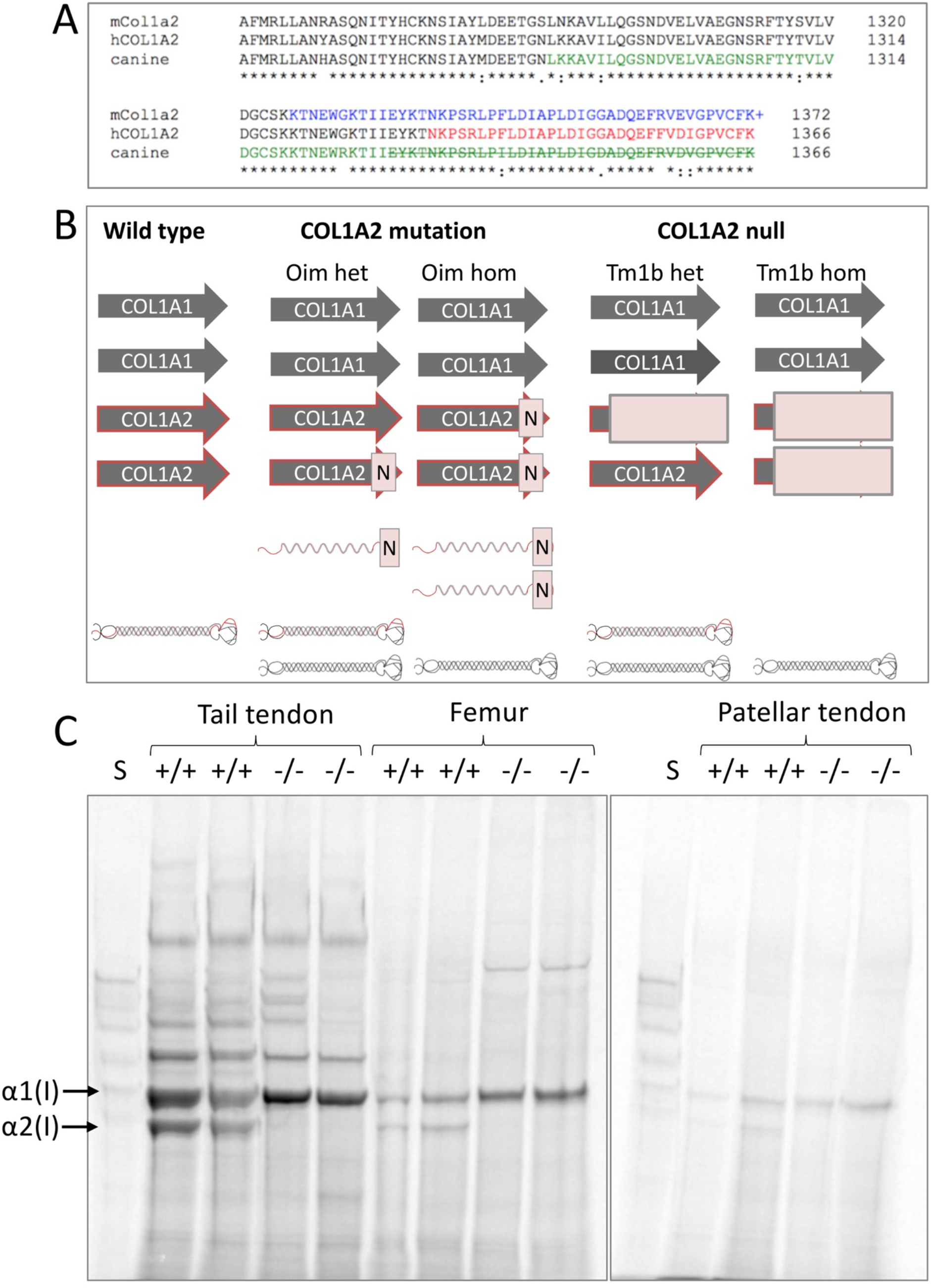
Col1a2 mutant and null alleles. A: Amino acid sequence alignment for the final 112 residues of the mouse (mCol1a2), human (hCOL1A2) and canine α2(I) collagen chains. Colour change indicates amino acid sequence changed from that shown due to frameshift caused by the oim or equivalent mutation. + indicates reported additional amino acid (18). Strike-through indicates truncation. B: Genetic differences between the oim and Col1a2 null lines with implications for collagen (I) protein synthesis. Arrows indicate COL1 genes; N indicates mutation, light red box indicates null allele. Folded heterotrimeric proteins are indicated in black and red, whilst homotrimers are in black only. The presence of unincorporated mutant Col1a2 allele is indicated as a red waveform with a mutation (N). C: Tendon and bone tissue from Tm1b wild-type (+/+) and homozygote (-/-) mice were labelled with [14C]proline and analysed by delayed reduction SDS-PAGE. No labelled α2(I) chain was present in tm1b homozygotes (-/-) unlike in wild-type controls (+/+) indicating that tm1b homozygotes are Col1a2 null. S; collagen standard. α1(I) and α2(I) chains are indicated.

The oim mutation is very similar to the first-identified mutation causing human osteogenesis imperfecta (autosomal recessive Silence type III), in which a 4 nucleotide deletion (c.4001_4004del) causes a frameshift (p.(Asn1334Serfs*34)) to alter the last 33 amino acids of the α2 chain of type I procollagen (Pihlajaniemi et al., 1984). The human mutation also results in homotrimeric type I collagen synthesis as the defective α2(I) chain cannot be incorporated into trimers (24, 25). In two additional patients with mild osteogenesis imperfecta, novel mutations were identified causing a 48 amino acid truncation of the α2 chain and a substitution of a cysteine residue important for interchain disulphide bonding respectively (26). In these cases the α2 chain was again synthesised but not incorporated into trimers. A similar mutation causing severe osteogenesis imperfecta in a Beagle dog was caused by a heterozygous 9 nucleotide replacement of a 4 nucleotide stretch, leading to a 37 residue truncation of the α2 chain and alteration of the final 44 amino acids of the truncated polypeptide (Campbell et al., 2001).

Oim heterozygotes do not show spontaneous fractures but appear to have a bone phenotype intermediate between that of wild-type and homozygous mice (20, 27–29). Similarly the parents of the human proband had no history or evidence of fractures but had a marked decrease in bone mass (30).

The osteogenesis imperfecta brittle bone phenotype, and the genetic association of collagen (I) homotrimer with osteoporosis (5, 31) contrasts with the phenotype of human patients in which mutations leading to nonsense-mediated decay of the COL1A2 mRNA cause a specific cardiac valvular form of Ehlers-Danlos syndrome (cv-EDS), not involving bone (32, 33) and evidence that Col1a2 silencing does not affect *in vitro* osteoblast mineralisation (34). EDS is generally characterised by hyperextensible skin and joint hypermobility. In cv-EDS patients the α2 chain of type I collagen is not synthesised, therefore all type I collagen molecules would be homotrimeric. It has been hypothesised that the phenotypic differences between the OI and cv-EDS patients could be explained by the cellular stress elicited by the presence of misfolded α2(I) procollagen chains in osteogenesis imperfecta (26, 35, 36), having a particularly detrimental effect on bone. Cellular stress has been implicated in human osteogenesis imperfecta caused by substitutions in the C-propeptide of the α-1(I) chain (37), in mouse models of triple-helical region mutations; Aga2 (90 a.a. extension to the α1(I) chain) (38), Brtl IV (G349C in α1(I)) (39) and Amish (G610C in α2(I)), identical to that found in a human kindred) (40), as well as in the zebrafish model Chihuahua (G574D in α1(I)) (41).

In this study we compared the bone phenotype of the oim model to that of a Col1a2 null mouse. We considered that comparing the oim model to that of a Col1a2 null line provided a unique opportunity to distinguish between the intracellular and extracellular effects of a collagen mutation linked to brittle bone disease, as well as to elucidate the effect of collagen (I) homotrimer on bone structure and mechanics. Specifically, we compared the bone phenotype of oim homozygous and heterozygous mutant mice with that of mice containing 1 or 2 copies of a targeted Col1a2 null allele and wild-type controls, in order to determine the contribution of collagen (I) homotrimer to bone fragility.

## Results

### Tm1b homozygotes lack the α2(I) collagen chain in bone and tendon

To verify lack of the α2(I) chain in tm1b homozygotes, tendon and bone samples from 8 week old mice were labelled with [14C]proline to detect newly synthesised collagen (I). SDS-PAGE analysis of labelled tissue extracts verified lack of α2(I) chain synthesis in tail tendon, bone and patellar tendon (Fig. 1C).

### Alterations to Mendelian inheritance in male mice of the oim and Col1a2 null lines

To determine if either the oim or Col1a2 alleles resulted in loss of mice prior to weaning, a chi-squared test was performed on genotype data for Col1a2 null and oim mice. There were significant differences between the observed and expected genotype percentages for male mice of both lines (oim; p=0.01, Col1a2 null; p=0.033), whereas no differences were seen for female mice of either line (Fig. 2 A–B). For oim males there was an 10.0% increase in wild-types, a 6.2% decrease in heterozygotes and a 3.8% decrease in homozygotes; whereas for Col1a2 null males there was an 8.0% increase in wild-types, a 9.6% decrease in heterozygotes and a 1.6% increase in homozygotes. Hence increased numbers of male wild-types and reduced numbers of male heterozygotes were observed for both lines. Conversely the data supported Mendelian inheritance in female mice.

**Figure 2.**
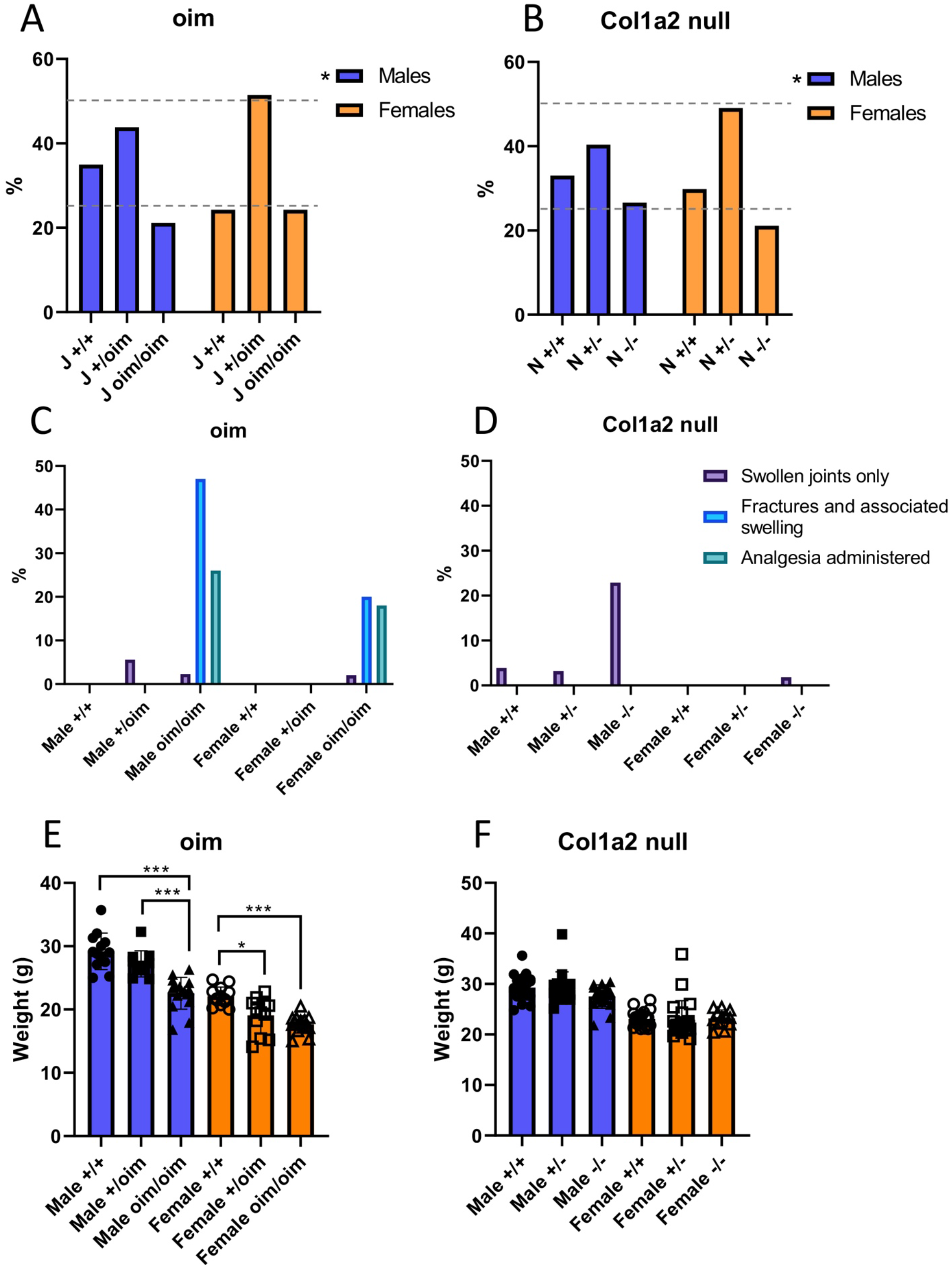
Inheritance pattern, musculoskeletal health summary and mouse weights for the oim and Col1a2 null lines. A-B: The percentage of mice of each genotype born to heterozygous parents for the oim (A) and Col1a2 null (B) lines. A chi-squared test showed significant differences between observed inheritance and expected Mendelian inheritance (25%, 50%, 25%; dashed grey lines) for male mice of both lines. C-D: The number of mice suffering from swollen joints and bone fractures as well as those treated with analgesics were recorded for the oim (C) and Col1a2 null (D) lines. These numbers are expressed as a percentage of total mouse numbers. A-D: n=71, 89 and 43 for male oim +/+, +/oim and oim/oim mice respectively; n=50, 106 and 50 for female oim +/+, +/oim and oim/oim mice respectively; n= 103, 126 and 83 for male Col1a2 null +/+, +/-, -/- mice respectively and; n=79, 130, 56 for female Col1a2 null +/+, +/-, -/- mice respectively. Swollen joints were mild in the Col1a2 null line and often not noted until advanced age, therefore the numbers shown above may be an underestimation due to many mice being culled at earlier time points. E-F: Weights of 18 week oim (E) and Col1a2 null (F) mice measured after euthanasia. Blue bars/filled shapes = males, orange bars/open shapes = females. * p-value < 0.05 and *** p-value <0.001. E: p-value for genotype <0.001 for males and females. E-F: n= 13, 11 and 16 for male oim male +/+, +/oim and oim/oim mice respectively, n=13, 12 and 15 for female oim +/+, +/oim and oim/oim mice respectively, n= 21, 17 and 17 for male Col1a2 null +/+, +/-, -/- mice respectively, n=21, 22, 12 for female Col1a2 null +/+, +/-, -/- mice respectively.

### Spontaneous fractures were observed solely in oim homozygotes, whilst male Col1a2 null homozygotes exhibited mild joint swelling

During colony maintenance it was noticeable that spontaneous fractures occurred in the oim line, but not in the Col1a2 null line. Mildly swollen ankle joints were however observed in male tm1b mice which was initially attributed to fighting. The proportion of mice with swollen joints, or noticeable fractures with swelling were determined, including those requiring analgesia (Fig. 2 C–D). Male mice demonstrated a more severe phenotype than female mice for both the Col1a2 null and oim lines. For the oim line fractures and associated swelling were observed in homozygous mice only, with an incidence of 47% for males (20 out of 43) and 20% (10 out of 50) for females. 26% of male mice (11 out of 43) and 18% of female mice (9 out of 50) received analgesic medication. Swollen joints alone presented in 6% of male oim heterozygotes (5 out of 89), 2% of male oim homozygotes (1 out of 43) and 2% of female oim homozygotes (1 out of 50). For the Col1a2 null line no bone fractures were observed and no analgesia was required to be administered. Swollen joints were observed predominantly in male homozygotes with an incidence of 23% (19 out of 83). Less than 4% of male wild-type (4 out of 103), heterozygous (4 out of 126) and female homozygous (1 out of 56) mice presented with swollen joints. No female wild-type or heterozygous mice were reported to have swollen joints. For mouse weights recorded after euthanasia at 18 weeks, male and female oim homozygotes were 22.7% and 20.0% lighter than wild-types (Fig. 2E). Heterozygotes were of intermediate weight with male homozygotes being 17.1% lighter than male heterozygotes and female heterozygotes being 13.6% lighter than wild-types. In contrast, there were no significant differences in weight between genotypes for the Col1a2 null line (Fig. 2F).

### Cortical bone analysis and three-point bending of bones from oim and Col1a2 null mice

Cortical thickness was significantly decreased in oim homozygotes; by 33% in 8 week males (Fig. 3A) and to a lesser extent at 18 weeks (22%) (Fig. 3F) and in females (14%; 8 wks, 20%; 18 wks). At 8 weeks cortical thickness in oim heterozygotes was intermediate between that of wild-types and oim homozygotes (a 20% reduction in males and 10% in females). At 18 weeks, but not significantly at 8 or 52 weeks, cortical thickness was also reduced in Col1a2 null homozygotes (18% in males and 10% in females). For oim homozygotes and heterozygotes polar moment of inertia was reduced at 8 weeks (Fig. 3B) in both males (65% and 42% respectively) and females (36% and 26% respectively), and at 18 weeks (Fig. 3G) in males (42% and 29% respectively), indicative of a reduced resistance to torsional loading. In contrast, this parameter was increased by 31% in male 8 week Col1a2 null homozygotes (Fig. 3B) as compared to their control (N +/+) (though not above the level of the oim wild-type control (J +/+)), and at 52 weeks in females by 30% as compared to heterozygotes (Fig. 3L). Periosteal circumference was decreased by 25% and 17% in male oim homozygotes and heterozygotes respectively at 8 weeks (Fig. 3C). However both periosteal (Fig. 3M) and endosteal circumference (Fig. 3N) increased in female Col1a2 null homozygotes at 52 weeks (by 9% and 16%), mirroring the same trend in males. Tissue mineral density was significantly higher in oim wild-types than the other genotypes at 8 weeks (Fig. 3E), but lower in male wild-types at 52 weeks (Fig. 3O).

**Figure 3.**
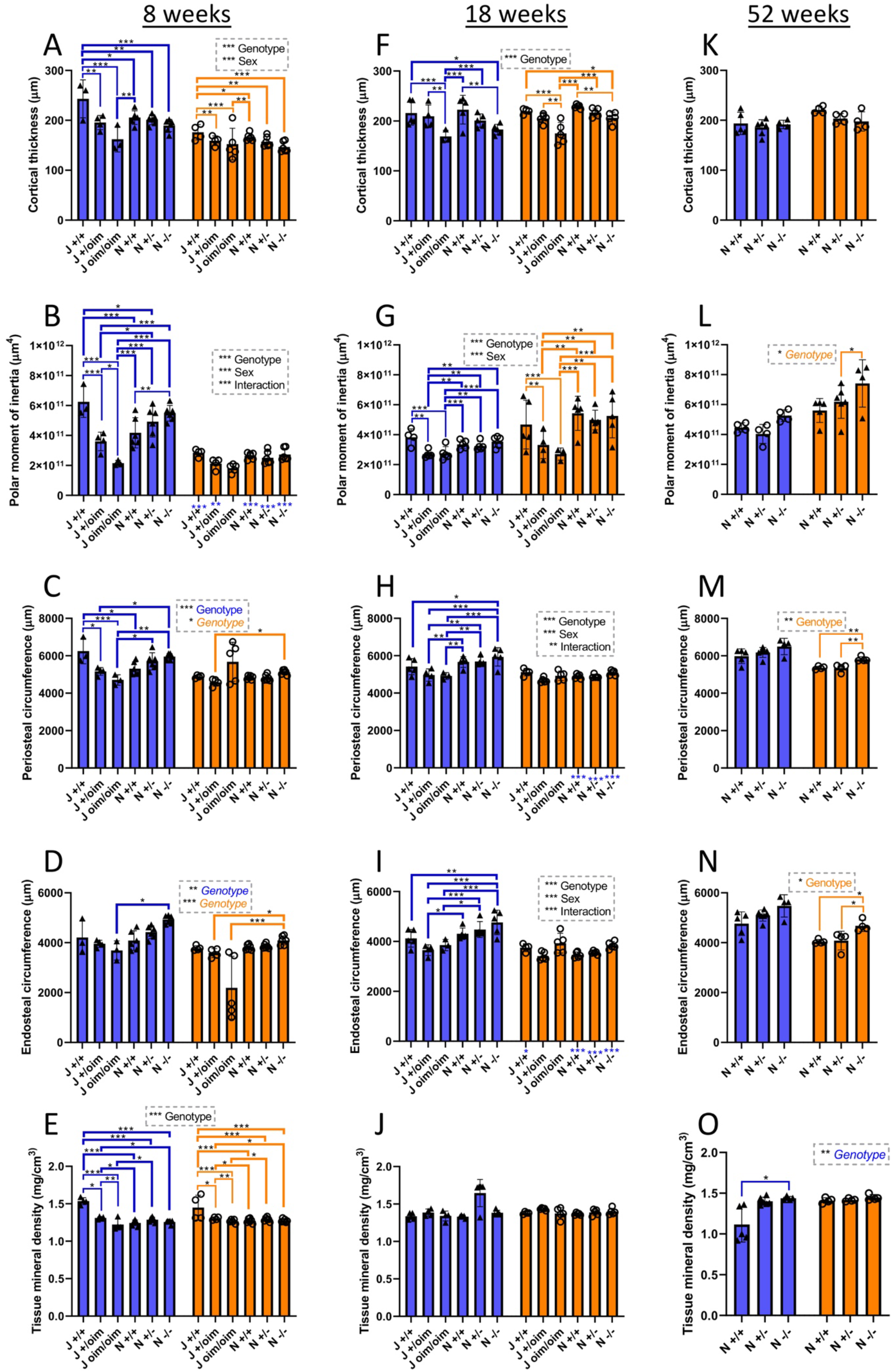
Femoral cortical bone analyses of oim and Col1a2 null mice. MicroCT scans were performed on the femur of oim and Col1a2 null mice at 8 (A-E), 18 (F-J) and 52 weeks (K-O). Reconstruction and analysis of scan files enabled determination of cortical thickness (A, F, K), polar moment of inertia (B, G, L), periosteal (C, H, M), and endosteal (D, I, N) circumference, as well as bone density (E, J, O). Blue bars/triangles = males, orange bars/circles = females. Thin blue (male) and orange (female) brackets show differences between genotypes within mouse lines and thick brackets between lines. Male/female differences for particular genotypes are shown below the x-axis for females (blue stars). * p-value < 0.05, ** p-value <0.01 and *** p-value <0.001. N=3 for male oim groups at 8 weeks, except for +/oim and bone density measurements where n=4. N=4 for female oim groups at 8 weeks, except for oim/oim where n=5. N=3 for male oim/oim at 18 weeks, except for bone density where n=4. N=4 for male oim/oim and female +/+, whilst n=5 for other oim groups at 18 weeks. N=6 for all Col1a2 null groups at 8 weeks and n=5 for all Col1a2 null groups at 18 weeks. N=4 for all Col1a2 null groups at 52 weeks, except male +/+ where n=5 and +/- where n=6.

To determine whether biomechanical properties of bones differed between the Col1a2 null and oim lines, femurs and tibias from both male and female mice at various ages were subjected to three-point bending to measure ultimate force, stiffness, stress and elastic modulus (Figs. 4–5). Ultimate force and stiffness were reduced in both male and female oim homozygotes at 8 and 18 weeks in the femur (Fig. 4 A, B, F, G) and the tibia (Fig. 5 A, B, F, G), although differences were not significant in 8 week tibia for which a non-parametric test was used. The cross-sectional area was also reduced at 18 weeks in the femur (by 15% and 22% in males and females respectively) (Fig. 4J), and at 8 weeks (44% in males, 18% in females) and 18 weeks (31% in males, 25% in females) in the tibia (Fig. 5 E, J), consistent with alterations to extrinsic biomechanical properties in oim homozygotes. Ultimate force reductions in the femur were 47% and 65% at 8 and 18 weeks respectively in males, and 51% at both ages in females (Fig. 4 A, F). In the tibia ultimate force was reduced by 54% in males and 44% in females at 18 weeks (Fig. 5F). Stiffness was reduced by 42% and 46% at 8 and 18 weeks in males and by 21% and 42% in females in the femur (Fig. 4B, G), and by 44% and 34% in males and females respectively in the tibia at 18 weeks (Fig. 5G). Only the 8 week femur showed a significantly lower ultimate force for oim heterozygotes (27% for males and 39% for females) (Fig. 4A) whilst stiffness was reduced only in female oim heterozygotes in the tibia at 18 weeks (by 27%) (Fig. 5G). The intrinsic biomechanical property, ultimate stress, was reduced in oim homozygotes in the femur at both 8 (51% in males, 43% in females) and 18 weeks (65% in males, 27% in females) (Fig. 4 C, H), but not in the tibia (Fig 5 C, H). Elastic modulus was unchanged in oim homozygotes in either the femur or tibia (Figs 4&5 D, I).

**Figure 4.**
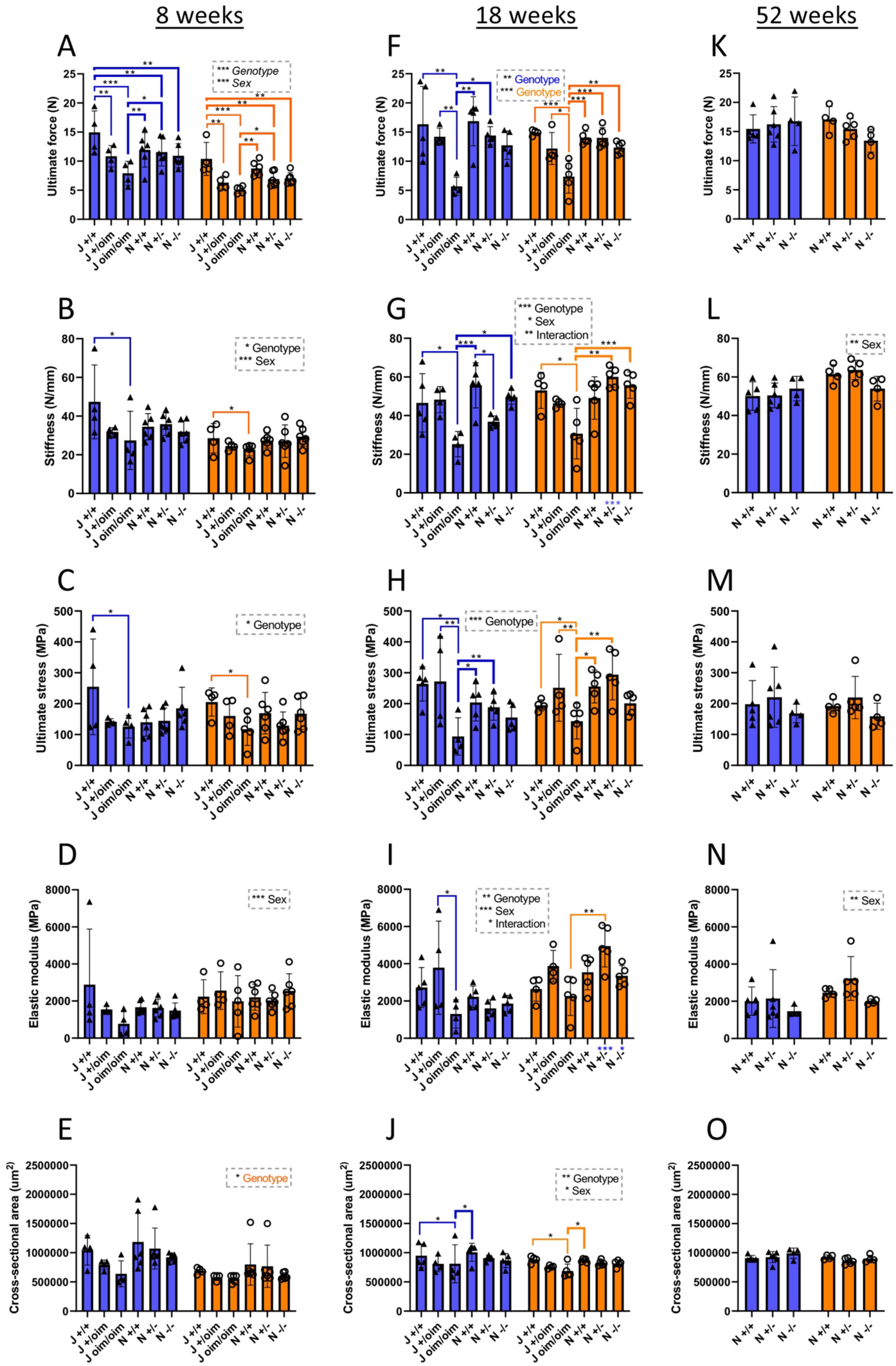
Three-point bending of femurs from oim and Col1a2 null mice. Femurs from oim and Col1a2 null mice were subjected to three-point bending at 8 (A-E), 18 (F-J) and 52 weeks (K-O). Ultimate force (A, F, K) and stiffness (B, G, L) (extrinsic) measurements were normalised to cross-sectional area (E, J, O) to calculate ultimate stress (C, H, M) and elastic modulus (D, I, N) (intrinsic). Blue bars/triangles = males, orange bars/circles = females. Thin blue (male) and orange (female) brackets show differences between genotypes within mouse lines and thick brackets between lines. Male/female differences for particular genotypes are shown below the x-axis for females (blue stars). * p-value < 0.05, ** p-value <0.01 and *** p-value <0.001. n=4 for all oim groups, except female J oim/oim and male 18 week J+/+ where n=5. n=6 for all Col1a2 null groups at 8 weeks and n=5 for all Col1a2 null groups at 18 weeks. n=4 for all Col1a2 null groups at 52 weeks, except female +/- and male +/+ where n=5 and male +/- where n=6.

**Figure 5.**
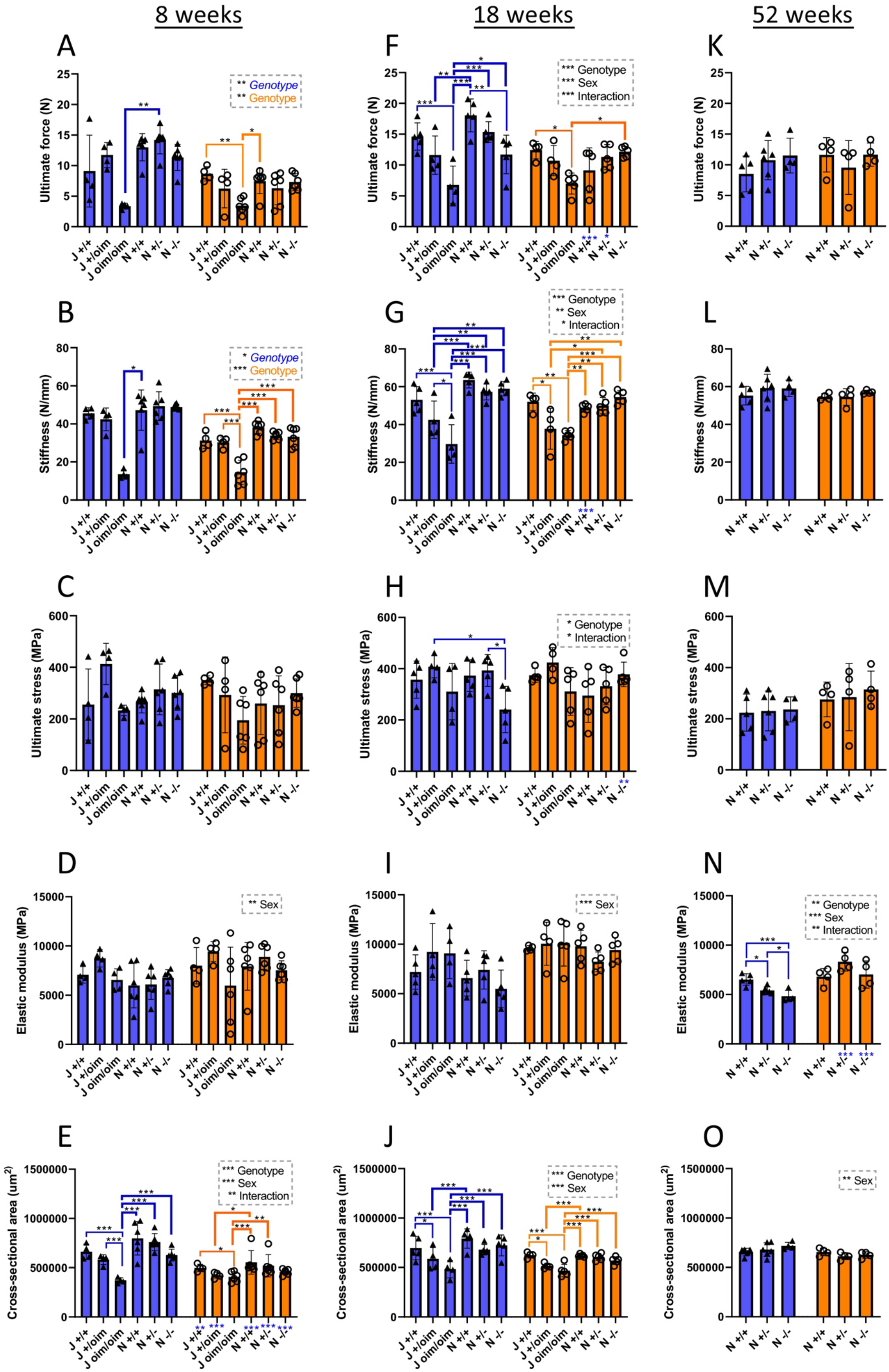
Three-point bending of tibias from oim and Col1a2 null mice. Tibias from oim and Col1a2 null mice were subjected to three-point bending at 8 (A-E), 18 (F-J) and 52 weeks (K-O). Ultimate force (A, F, K) and stiffness (B, G, L) (extrinsic) measurements were normalised to cross-sectional area (E, J, O) to calculate ultimate stress (C, H, M) and elastic modulus (D, I, N) (intrinsic). Blue bars/triangles = males, orange bars/circles = females. Thin blue (male) and orange (female) brackets show differences between genotypes within mouse lines and thick brackets between lines. Male/female differences for particular genotypes are shown below the x-axis for females (blue stars). * p-value < 0.05, ** p-value <0.01 and *** p-value <0.001. n=4 for all oim groups, except female 18 week oim/oim and male 18 week J +/+ where n=5, and female 8 week oim/oim where n=6. n=6 for all Col1a2 null groups at 8 weeks and n=5 for all Col1a2 null groups at 18 weeks. n=4 for all Col1a2 null groups at 52 weeks, except male +/+ where n=5 and male +/- where n=6.

Deletion of Col1a2 had no effect on 3-point bending parameters in the femur (Fig. 4). In the tibia only ultimate force was reduced (by 35%) in males at 18 weeks of age as compared to the wild-type controls (Fig. 5F), whilst ultimate stress was reduced by 39% as compared to Col1a2 null heterozygotes (Fig. 5H). At 52 weeks only the elastic modulus was affected in males, being reduced by 26% in Col1a2 null homozygotes and 17% in heterozygotes.

### Analysis of trabecular bone of oim and Col1a2 null mice

To analyse differences in trabecular bone structure between genotypes, the proximal tibias from oim and Col1a2 null mice were analysed by μCT (Fig. 6).

**Figure 6.**
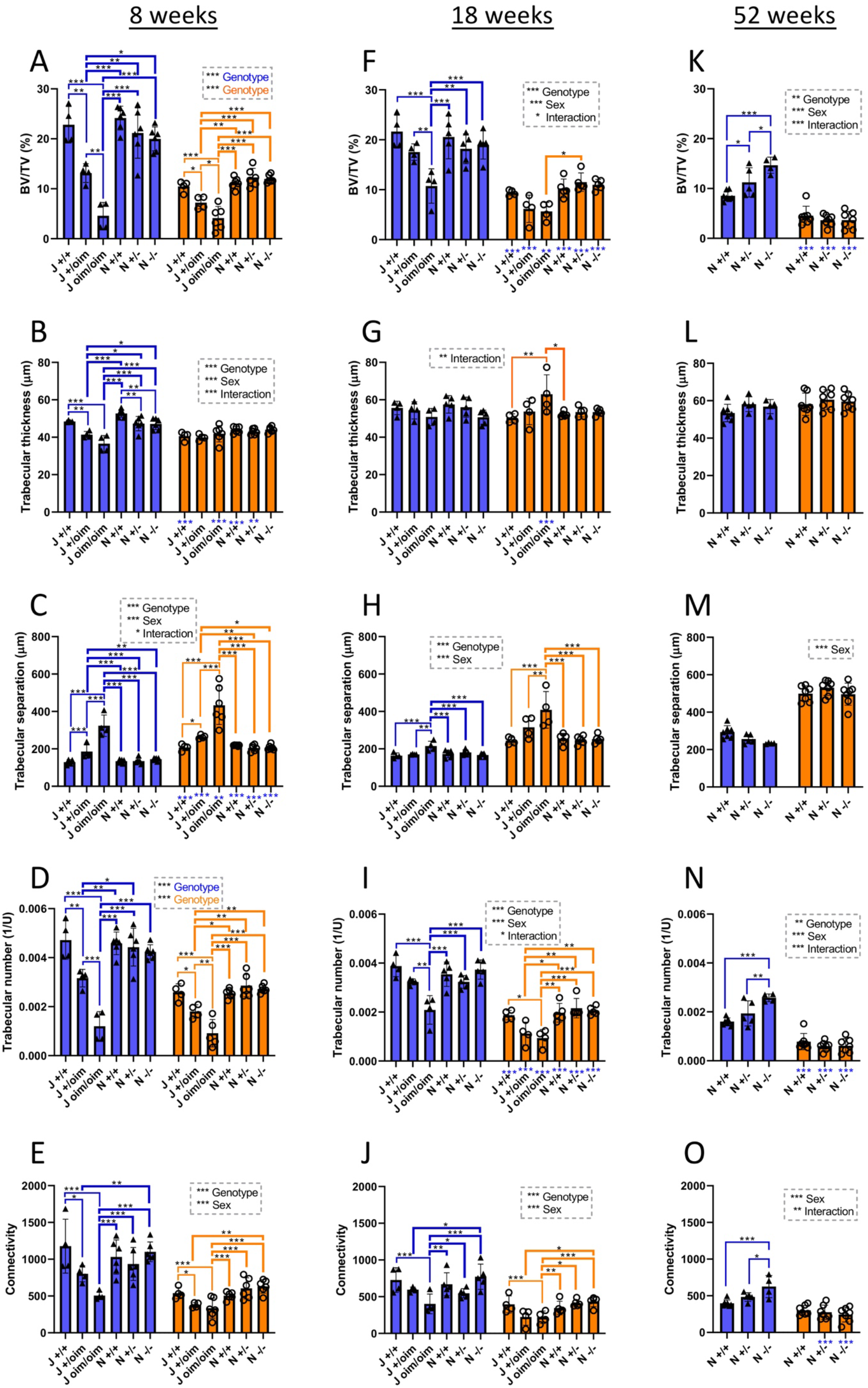
Micro-computed tomography bone scans of oim and Col1a2 null mice. MicroCT scans were performed on the knee joints of oim and Col1a2 null mice at 8 (A-E), 18 (F-J) and 52 weeks (K-O). Reconstruction and analysis of scan files enabled determination of bone volume (BV/TV) (A, F, K), trabecular thickness (B, G, L), trabecular separation (C, H, M), trabecular number (D, I, N) and connectivity (E, J, O). Blue bars/triangles = males, orange bars/circles = females. Thin blue (male) and orange (female) brackets show differences between genotypes within mouse lines and thick brackets between lines. Male/female differences for particular genotypes are shown below the x-axis for females (blue stars). * p-value < 0.05, ** p-value <0.01 and *** p-value <0.001. n=4 for all oim groups, except female 8 week old oim/oim where n=6 but n=5 in D. n=6 for all Col1a2 null groups at 8 weeks and n=5 for all Col1a2 null groups at 18 weeks. n=7 for all Col1a2 null groups at 52 weeks, except male +/- where n=5 and male -/- where n=4.

There were significant changes in bone volume and architecture in both males and females carrying the oim mutation at both 8 weeks and 18 weeks of age, and most of the differences showed an apparent gene dose effect. In males at 8 weeks, BV/TV was decreased by 42% in heterozygote mice and 80% in homozygous mice compared to wild-type controls (Fig. 6A), due to a decrease in both trabecular thickness (Fig. 6B) and trabecular number (Fig. 6D). The decrease in trabecular thickness and number led to a 151% increase of trabecular separation in oim/oim homozygotes (Fig.6C). Similar changes were observed in female mice at 8 weeks and whilst still pronounced the differences tended to be smaller, and trabecular thickness was unaffected (Fig. 6B). Differences observed for heterozygotes of both sexes were less than those of the respective homozygotes.

At 18 weeks of age, the pattern of the effects of the oim mutation on trabecular bone were similar to those observed at 8 weeks, however the differences between wild-types, heterozygotes and homozygotes were generally smaller. For instance, BV/TV in males was decreased by 50% at 18 weeks (Fig. 6F), compared to 80% at 8 weeks of age (Fig. 6A). There was no longer a significant reduction in trabecular thickness in males, and even an increase in female homozygotes (Fig. 6G).

The only similar alterations resulting from Col1a2 deletion were an 11% decrease in trabecular thickness in 8 week old male null homozygotes and a 10% decrease in heterozygotes (Fig. 6B). Conversely at 52 weeks in males there were Col1a2 loss-dependent increases in BV/TV and trabecular number, with increased connectivity in homozygotes but not heterozygotes. No differences were detected between wild-types of each strain, indicating that this had no influence on the bone structural parameters measured.

### Age-related deterioration of Col1a2 null male homozygotes

The tm1b line (Col1a2 null) was maintained on a mild protocol and mice exceeding the mild severity limit were legally required to be humanely killed. We noted an unexpectedly high loss of male Col1a2 null homozygotes due to a loss of condition, including weight loss and respiratory difficulties. A Kaplan-Meier ‘survival’ analysis was performed on the Col1a2 null mouse line (Fig. 7). All genotyped mice from this line were included in the analysis up to the age of 12 months, the end point for all experiments. The majority of genotypes had very few losses throughout the time course, with no animals lost for female wild-types and heterozygotes and only one animal lost for female homozygotes, male wild-types and heterozygotes out of a total of 19 mice. In contrast, almost 50% of male homozygotes were lost over the 12 month experimental period.

**Figure 7.**
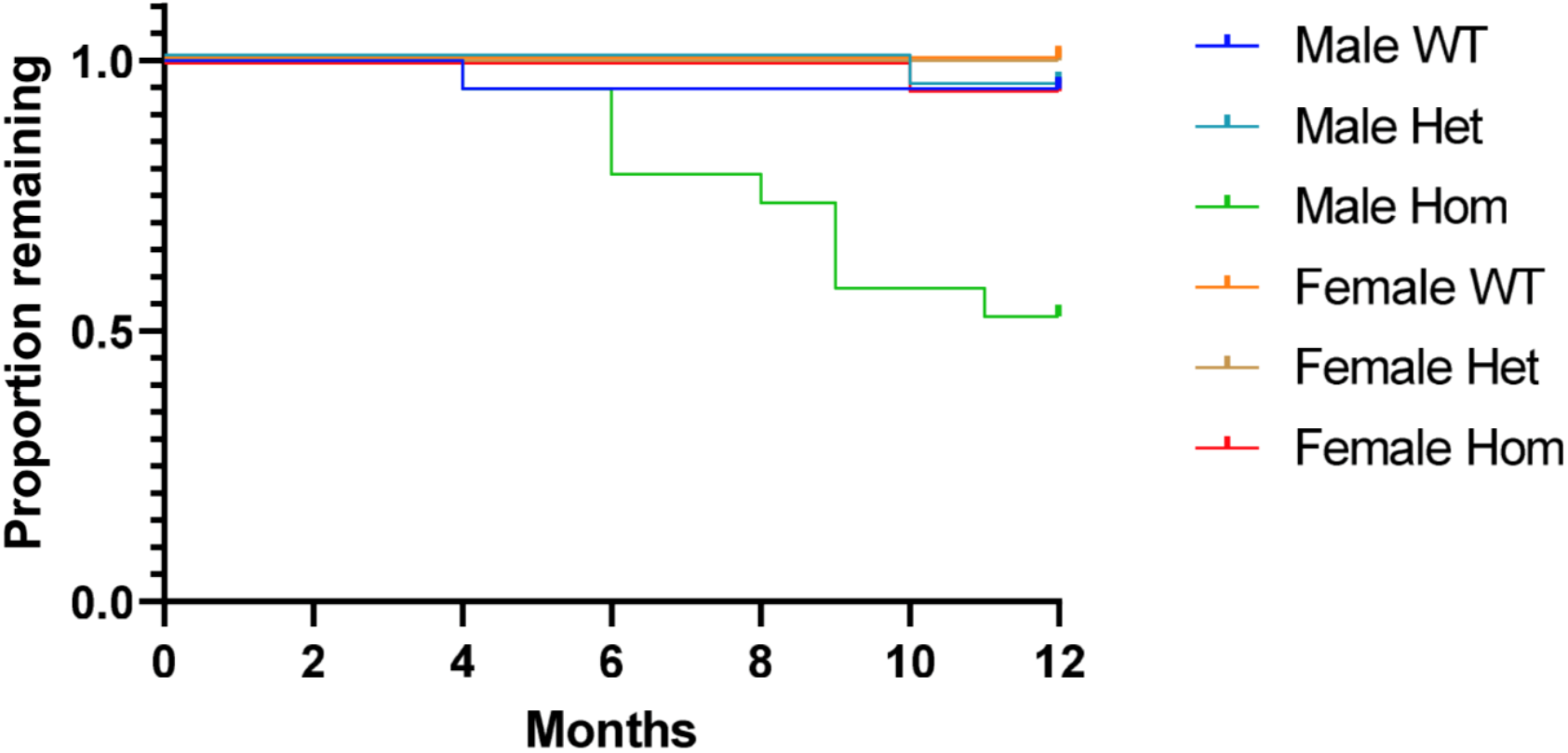
Survival analysis for the Col1a2KO mouse line. A Kaplan-Meier ‘survival’ analysis was performed on all genotyped mice from the Col1a2 null line up until 12 months (end of experiment). A log-rank (Mantel-Cox) test was performed which gave an overall p-value of <0.0001 indicating a significant difference between survival curves. n=19 for all groups.

### Oim heterozygotes do not down-regulate the mutant allele

We considered that the less severe bone phenotype of oim heterozygotes could be related to a compensatory down-regulation of mRNA from the oim mutant allele. A custom allelic discrimination assay indicated that mRNA from both alleles was present in bone tissue from heterozygotes at both 8 (Fig. 8A) and 18 weeks of age (Fig. 8B). Whilst other tissues can be affected in the oim line the bone phenotype is particularly severe. We therefore determined whether there was compensatory downregulation of the mutant allele in tendon at 8 weeks (Fig. 8C) and 18 weeks (Fig. 8D), and in aorta (Fig. 8E), kidney (Fig. 8F), liver (Fig. 8G), or lung (Fig. 8H) at 18 weeks. For all tissues examined, mRNA from both alleles was present at a close to 50-50% ratio in heterozygotes.

**Figure 8.**
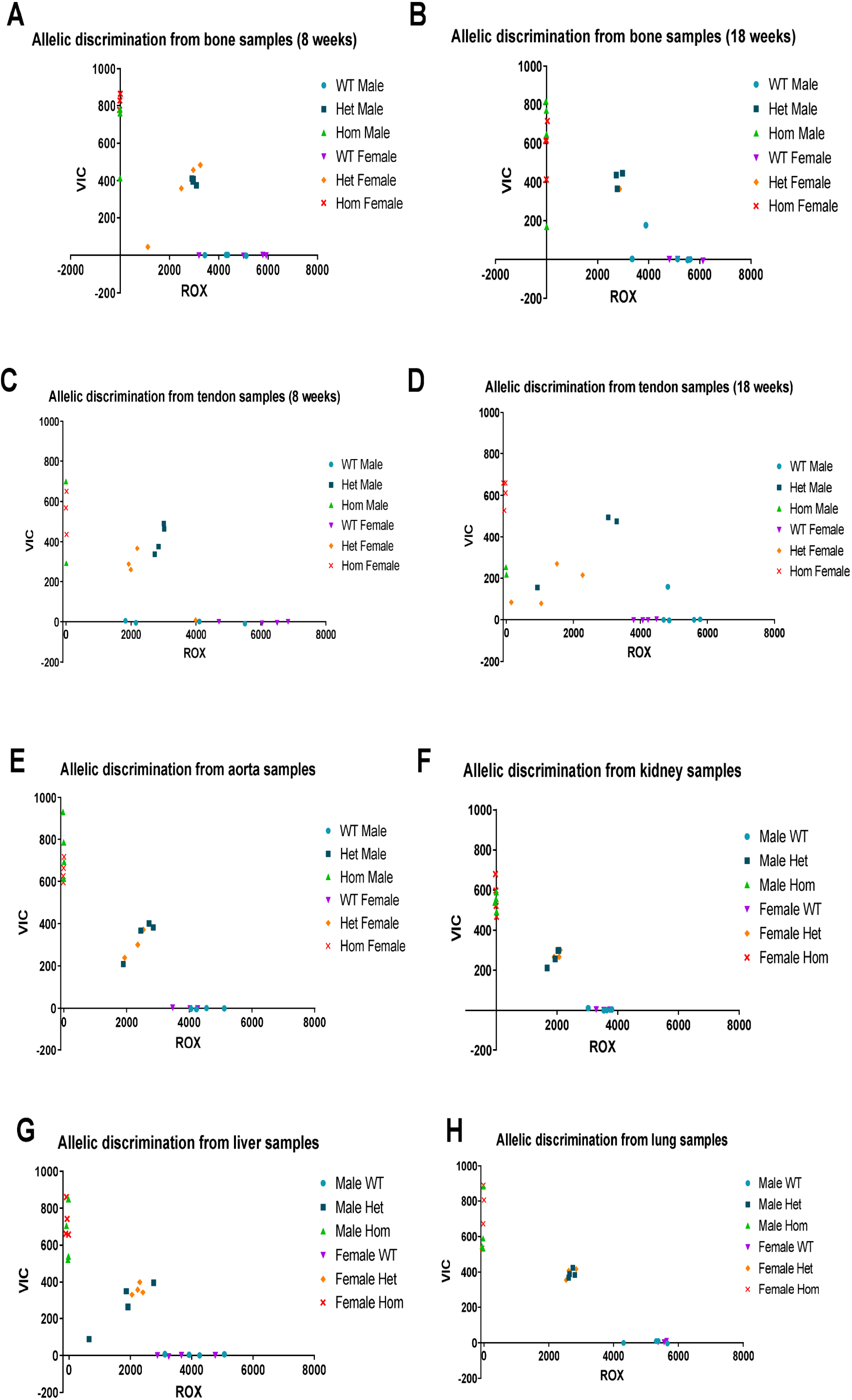
Allelic discrimination in the oim line. Allelic discrimination was carried out in bone (A-B) and tendon (C-D) at 8 (A, C) and 18 weeks (B, D) and in aorta (E), kidney (F), liver (G) and lung (H) at 18 weeks using ROX labelled primer/probes for the wild-type and VIC for the oim allele channels. For all tissues wild-types displayed little to no VIC signal, whilst homozygotes displayed no ROX signal. Heterozygotes had approximately intermediate signals in both channels with no systematic deviations between sexes or tissues. n=4 for all groups except 8 week bone and tendon samples from male and female oim/oim where n=2-3, 18 week bone and tendon samples from male +/+ where n=5 and female +/oim where n=1, kidney, aorta and lung samples for female +/+ and +/oim where n=3.

### There is a genetic interaction between the oim mutant allele and collagen (I) homotrimer

As the mutant allele is not downregulated in oim heterozygotes, it is feasible that the less severe phenotype relates to gene dosage; given that heterozygotes have 1 rather than 2 copies of the mutant allele. To test this hypothesis we crossed the oim and tm1b lines to produce compound heterozygote offspring, along with heterozygotes of each genotype and wild-type controls (Fig. 9A). Compound heterozygotes contain only one copy of the oim mutant allele but have no wild-type Col1a2 allele so produce solely homotrimeric α1 type I collagen.

**Figure 9.**
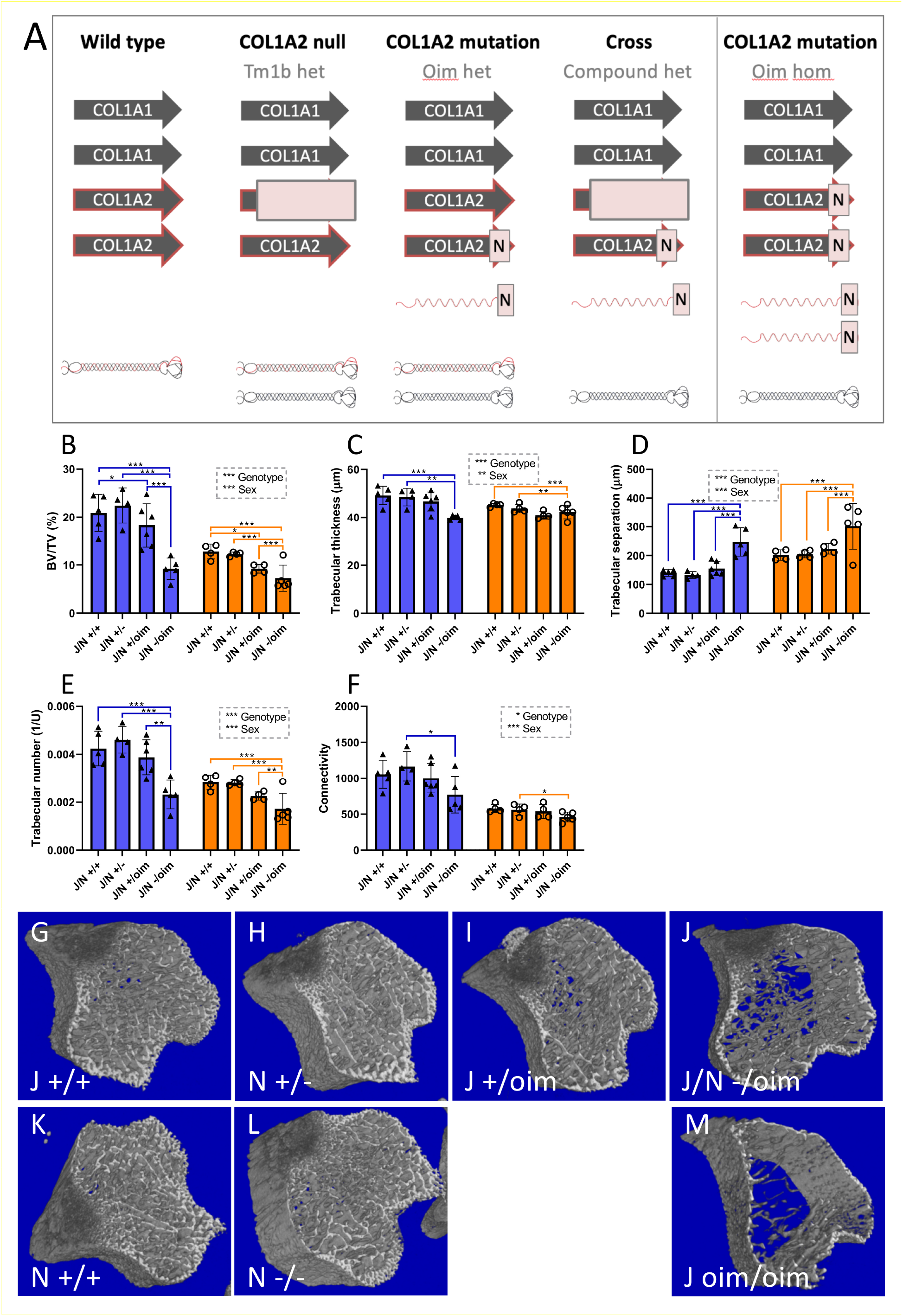
Bone structural properties are impaired in compound heterozygotes as compared to heterozygotes of the oim or Col1a2 null lines. A: Genetic differences between the heterozygous oim and Col1a2 null alleles and the compound heterozygous allele, with implications for collagen (I) protein synthesis. The homozygous oim allele is shown for comparison. Arrows indicate COL1 genes; N indicates mutation, light red box indicates null allele. Folded heterotrimeric proteins are indicated in black and red, whilst homotrimers are in black only. The presence of unincorporated mutant Col1a2 allele is indicated as a red waveform with a mutation (N). B-F: MicroCT scans were performed on the knee joints of offspring from heterozygous crosses of each line. Reconstruction and analysis of scan files enabled determination of bone volume (B), trabecular thickness (C), trabecular separation (D), trabecular number (E) and connectivity (F). Blue bars/triangles = males, orange bars/circles = females. Blue and orange brackets show differences between genotypes within males (blue) or females (orange). * p-value < 0.05, ** p-value <0.01 and *** p-value <0.001. n=4 for all groups, except male J/N +/+ and J/N - /oim of both sexes where n=5, and male J/N +/oim where n=6. G-M: Representative scan images from wild-types from the oim line (G), Col1a2 null heterozygotes (H), oim heterozygotes (I), compound heterozygotes (J), wild-types from the Col1a2 null line (K), Col1a2 null homozygotes (L) and oim homozygotes (M). Bone structural defects are more pronounced in compound than in oim heterozygotes.

The proximal tibias from 8-week-old oim and Col1a2 null (oim/Col1a2 null) cross line mice were analysed by μCT (Fig. 9 B–F). Oim/Col1a2 null compound heterozygotes demonstrated a significantly reduced bone volume (by 49% in males and 21% in females) (Fig. 9B), increased trabecular separation (by 60% in males, 35% in females) (Fig. 9D) and reduced trabecular number (by 40% in males and 24% in females) (Fig. 9E) compared to oim heterozygotes. No significant differences in trabecular thickness (Fig. 9C) and connectivity (Fig. 9F) were detected between compound heterozygotes (-/oim) and oim heterozygotes (+/oim) though differences between Col1a2 null heterozygotes (+/-) and compound heterozygotes were present for connectivity and trabecular thickness, and also between compound heterozygotes and wild-types (+/+) for trabecular thickness. Representative scan images (Fig. 9 G’M) support a structural deterioration with the oim allele as (+/oim) > (-/oim) > (oim/oim), indicating that gene dosage alone does not determine phenotypic severity.

The cortical bone of the femur was also analysed as an indicator of biomechanical properties. Cortical thickness was reduced by 28% in males and 13% in female compound heterozygotes as compared to oim heterozygotes (Fig. S1A). Differences were not detected for other cortical bone parameters (Fig. S1 B–E). Notably in the main 8 week cohort (Fig. 3) differences in cortical bone parameters were only detected between oim heterozygotes and homozygotes for polar moment of inertia in males and for tissue mineral density.

The data for the compound heterozygotes was compared to that of the oim homozygotes and the Col1a2 null line, all of which produce no heterotrimeric type I collagen (Fig. S2). Oim homozygotes and compound heterozygotes demonstrated a reduced bone volume (Fig. S2A) and trabecular thickness (males only) (Fig. S2B), increased trabecular separation (Fig. S2C), and reduced trabecular number (Fig. S2D) and connectivity (Fig. S1E) as compared to Col1a2 null homozygotes. For trabecular parameters, excepting trabecular thickness (Fig. S2B), the bone structural properties of oim homozygotes were inferior to that of compound homozygotes, which were themselves inferior to that of Col1a2 null homozygotes. For example bone volume (Fig. S2A) was reduced by 54% in male and 42% in female compound heterozygotes, but by 77% in male and 67% in female oim homozygotes as compared to Col1a2 null homozygotes. For cortical parameters, cortical thickness was lower in compound heterozygotes than in oim homozygotes or Col1a2 null homozygotes (Fig. S2F), whilst polar moment of inertia (Fig. S2G), periosteal circumference (males only) (Fig. S2H) and endosteal circumference (Fig. S2I) were lower in oim homozygotes and compound heterozygotes than in Col1a2 null homozygotes. Endosteal circumference mirrored the effect on most trabecular parameters with oim homozygotes having inferior bone structural properties than compound heterozygotes, whilst for polar moment of inertia and periosteal circumference (as with trabecular thickness in males) the oim heterozygotes and compound heterozygotes were comparable. Cortical thickness was unusual in that compound heterozygotes had inferior properties to oim homozygotes. Therefore the bone phenotype of the compound heterozygotes was considerably more severe than the Col1a2 null lacking any mutant α2(I) chain, but generally less severe than or similar to the oim homozygotes with two mutant alleles.

## Discussion

The α2 chain of type I collagen appears largely indispensable to vertebrate life, with heterotrimeric ((α1)_2_(α2)_1_) type I collagen predominating in tetrapods. Actinopterygians (ray-finned fish) even contain a third alpha chain (α3) (42) and there is no obvious homotrimeric (α13) type I collagen precursor in more distantly related organisms (4, 43). Human and mouse α2(I) chain mutations resulting in homotrimeric type 1 collagen were known to cause brittle bones (18, 25), whilst humans with COL1A2 null alleles have cardiac valvular EDS but no overt bone phenotype (32). However common COL1A1 alleles resulting in homotrimeric type I collagen are associated with osteoporosis (5). Here we have shown that homotrimeric collagen *per se* largely does not affect bone structural and mechanical properties, but there is pre-weaning loss of male heterozygotes and age-related loss of male null homozygotes as well as a detrimental genetic interaction between the oim mutant Col1a2 allele and the homotrimeric form.

To determine the contribution of collagen (I) homotrimer to bone fragility, we compared the bone phenotype of two mouse lines that lack the α2 chain of type I collagen: the oim mutant, a Col1a2 null and a combination of the lines. After propagating the lines, we measured a difference in Mendelian inheritance for males from both lines; interestingly this seemed to increase the proportion of wild-type males whilst decreasing the proportion of heterozygotes. To our knowledge pre-weaning loss of male heterozygotes has not been reported for either line. Enzymatic susceptibility assays and differential scanning calorimetry have previously indicated that tail tendon from heterozygous oim mice contains both heterotrimeric and homotrimeric type I collagen (44, 45). Reconstituted fibrils comprising both heterotrimeric and homotrimeric collagen (I) molecules showed subfibrillar segregation of each trimeric form (46). Hence, mixed fibrils in heterozygotes may affect tissue remodelling or mechanics during development resulting in decreased survival of heterozygotes.

Unlike oim homozygotes, we observed no fractures in Col1a2 null homozygotes. Bone structural parameters and material properties were largely similar between wild-type, heterozygote and Col1a2 null homozygotes. However, there was a reduction in femoral cortical thickness at 18 weeks, and for males a reduction in ultimate force in the tibia at 18 weeks and trabecular thickness at 8 weeks; paralleling the trend but not the extent of the reduction observed in oim homozygotes. Our results for the oim line at 8 weeks are in agreement with previous studies showing a significant reduction in tibial bone volume and trabecular thickness in oim homozygotes (47) and a significant reduction in ultimate force, stiffness and ultimate stress of the femur (48). By including sex as a factor in our analyses we noted the decrease in tibial trabecular thickness was particular to male homozygotes. Our results for the oim line at 18 weeks were similar to those previously reported for 4 month old mice on the same background although with some differences relating to the significance or gender/sex dependence of the differences (29). In accordance with previous studies, we found no difference in intrinsic elastic modulus or ultimate stress of the tibia of oim homozygotes compared to wild-type controls (29, 48, 49), although ultimate stress but not elastic modulus was decreased in the femur. Decreased femoral ultimate stress was previously reported for 12-14 week oim homozygotes (50, 51), but not in all studies (52). Oim homozygote mice were lighter and visually smaller than their wild-type and heterozygote littermates, therefore the differences seen in extrinsic but not intrinsic mechanical properties, particularly of the tibia, between oim homozygotes and wild-types could imply that the increased bone fragility of oim homozygotes is due to the reduced size of these mice. Cross-sectional area was significantly reduced in the tibia at both ages, and in the femur at 18 weeks, though not at 8 weeks. microCT analysis however indicates some intrinsic differences in bone structure, with reduced cortical and trabecular bone.

Col1a2 null homozygotes did not display bone fragility but swollen joints were identified in males who also displayed age-related deterioration in condition. Human COL1A2 null homozygotes have cardiac valvular Ehlers-Danlos syndrome (cvEDS) (32, 33) hence the age-related deterioration in mice may relate to cardiovascular abnormalities. Indeed the International Mouse Phenotyping Consortium reports dilated left heart ventricle in males measured at 12 weeks of age and increased heart weight at 16 weeks in Col1a2 null homozygotes (53). Cardiovascular abnormalities were however observed in mice of both sexes and cvEDS patients are not solely male. It may be that cardiovascular defects present earlier in male mice due to increased activity or remain subclinical in females. IMPC also reported a skeletal phenotype for Col1a2 null male mice with increased (notably not decreased) bone mineral content and density (DEXA, 14 weeks) and “abnormal” femur and tibia morphology (X-ray, 14 weeks). Reported phenotypes change over time due to continual addition of control samples, but alterations to body fat in males and increased circulating alkaline phosphates in females were also listed. In the present study, loss of Col1a2 appeared to improve some bone structural parameters by 52 weeks (periosteal and endosteal circumference in females, tissue mineral density, trabecular bone volume, trabecular number and connectivity in males), possibly reflecting lower turnover of homotrimeric collagen (I). Elastic modulus was however reduced in Col1a2 null males by 52 weeks indicative of some altered bone material properties. Overall a combination of early loss of heterozygous males and age-related deterioration in null homozygotes could exert selective pressure to maintain the heterotrimeric form of type I collagen in vertebrate populations.

Whilst the strain of the lines differs slightly (C57BL/6J versus C57BL/6N), similar cortical and trabecular bone parameters have been reported in both strains (54). In the present study, differences were detected between wild-types of each strain at 8 weeks for cortical thickness (Fig. 3A) and tissue mineral density (Fig. 3E) in both sexes, and for polar moment of inertia (Fig. 3B) and periosteal circumference (Fig. 3C) in males. However for cortical thickness and polar moment of inertia the oim homozygotes (J oim/oim) were still significantly lower than the Col1a2 wild-types (N +/+) as well as than wild-types of their background strain (J +/+). For 3-point bending parameters of either the femur or tibia there were no significant differences between wild-types of either strain. Hence solely periosteal circumference and tissue mineral density differences may be influenced by the background strain. Reduced periosteal circumference in male compound heterozygotes (but not cortical thickness, polar moment of inertia or endosteal circumference) could therefore be influenced by the mixed 6J/6N background.

A key finding of this study was that the bone phenotype of a single copy of the oim allele was exacerbated by the absence of heterotrimeric type I collagen; i.e. that oim heterozygotes had a less severe phenotype than compound oim/null heterozygotes. Notably the phenotype of oim homozygotes still appeared more severe than that of compound heterozygotes, indicating some gene dosage effect for the mutant allele. Oim heterozygotes do not down-regulate the mutant allele (Fig. 7) hence presumably there is no mechanism to detect the oim mutation prior to translation or trimerization. Despite the severe bone phenotype in oim homozygotes, and whilst ER stress has been demonstrated in other OI models (37–41) we detected no evidence of increased ER stress by qPCR or by osteoblast ultrastructural analysis (not shown). A report indicating that relieving ER stress improves femoral mechanical properties in oim heterozygotes, actually demonstrated no difference between placebo and treatment groups and did not monitor ER stress (55). We did observe several significant differences between oim heterozygotes and wild-type controls, including all tibial structural parameters and femoral ultimate force at 8 weeks. At 18 weeks differences were limited to tibial stiffness in females and tibial cross-sectional area, consistent with a previous study which reported few significant differences between wild-types and heterozygotes at 18 weeks (Yao et al., 2013). Hence bone structural parameters can improve between 8 and 18 weeks in both heterozygotes and homozygotes. In oim heterozygotes at 8 weeks it is unclear if the observed differences in bone parameters relate solely to the interaction between the oim mutant allele and the proportion of homotrimeric collagen (I) that was present, or if the oim allele alone exerts an effect. The genetic interaction between the oim allele and homotrimeric collagen (I) could relate to the process of collagen folding and trimerization within the endoplasmic reticulum. The observed allelic series is consistent with a model in which the abnormal alpha-2 chain trimerises with two normal alpha-1 chains, but results in trimer degradation and concurrent suppression of collagen fibril assembly. Alternatively the homotrimeric collagen (I) could alter cell-matrix interactions to modulate cellular stress responses, if present. Procollagen N- and C-propeptides, derived from the proteolytic cleavage of procollagen, have been shown to have intracellular roles in modulating protein synthesis (56–58) whilst homotrimeric triple helical regions increased proliferation and migration as compared to heterotrimeric pepsinised collagen (11). Hence homotrimeric forms of collagen fibrils or propeptides have the potential for altered signalling affecting cellular stress responses. The association of homotrimeric type I collagen with several common human age-related diseases, in which it is unlikely to predominate structurally, could therefore relate to altered cellular signalling or cell stress responses.

The experiments outlined above demonstrate a genetic interaction between homotrimeric collagen (I) and the oim mutant allele, suggesting that the presence of heterotrimeric collagen (I) in oim heterozygotes alleviates the effect of the oim mutant allele in bone.

## Materials and Methods

### Mouse models

A Col1a2 knock-out mouse line (Col1a2^tm1b(EUCOMM)Wtsi^, C57BL/6N) (Col1a2 null, N) and osteogenesis imperfecta mouse line (Col1a2^oim^, C57BL/6J) (oim, J) were used to investigate the effects of the absence and the mis-folding of the α2(I) polypeptide chain respectively (Fig. 1B). The Col1a2^tm1b(EUCOMM)Wtsi^ line was derived from Col1a2^tm1a(EUCOMM)Wtsi^, purchased from the Mutant Mouse Resource and Research Centre (MMRRC) at UC Davies, by Cre-mediated recombination (59) during IVF provided by MRC Harwell. The Col1a2^oim^ line was a kind gift from Prof. Charlotte Phillips, University of Missouri and was subsequently rederived using Charles River (Massachusetts, USA) services. The strains have been deposited and are available from the MMRRC (RRID 66964, C57BL/6N-Col1a2<tm1b(EUCOMM)Wtsi>/LaelMmnc and RRID 66518, B6J.Cg-*Col1a2^oim^*/McbrMmnc). Mice were housed at the University of Liverpool in a specific pathogen free unit in groups of up to 5 by litter, with oim homozygotes and Col1a2 null/oim heterozygotes housed separately after weaning. Food and water were supplied *ad libitum* and wet food was supplied to oim homozygotes and Col1a2 null/oim heterozygotes due to fragile teeth. Cage balconies were removed for oim homozygotes and Col1a2 null/oim heterozygotes to reduce fracture risk and non-tangling bedding was supplied as standard for all mice. The mice were housed in the same room at 20-24°C and 45-65% humidity with a 12 hour light/dark cycle. All breeding and maintenance of animals was performed under project licences PP4874760 and P92F55CB2, in compliance with the Animals (Scientific Procedures) Act 1986 and UK Home Office guidelines. Details of all animals were recorded on tick@lab (a-tune) laboratory animal management software (Darmstadt, Germany), including health and treatment reports. Genotyping was carried out using Transnetyx (Tennessee, USA) services using the ‘Col1a2-2 WT’ probe with ‘LAC Z’ or ‘L1L2-Bact-P TA’ for tm1a allele or ‘L1L2 tm1b’ for tm1b allele, and the ‘oim’ probe for the oim line. Blinding was carried out by processing mice and labelling samples according to the mouse number, rather than genotype, however KJL was responsible for the initial allocation of animals to specific experimental groups. Wild-type (N +/+, J +/+), heterozygote (N +/-, J +/oim) and homozygote (N -/-, J oim/oim) mice were sacrificed at 8 (±3 days) and 18 (±3 days) weeks for analysis as well as 52 weeks (±8 days) for the Col1a2 null line. Eight and 18 weeks were chosen as the time points at which long bone growth and bone mineralisation respectively are complete. Oim mice were not maintained up to 52 weeks due to welfare considerations for homozygotes exhibiting spontaneous fractures. Cross-breeding of both lines was also performed (Col1a2 null/oim, mixed background) and wild-type (+/+), Col1a2 null heterozygote (+/oim), oim heterozygote (+/-) and compound heterozygote (-/oim) mice were sacrificed at 8 weeks. A total of 281 mice were used in this study, excluding those from which solely ‘survival’ and health analyses were derived. No mice were excluded from experimental groups.

### Mouse dissection

Mice were culled using a rising carbon dioxide concentration method in an automated CO2 delivery chamber. After confirmation of the permanent cessation of the circulation, the mice were weighed and the tail was removed at the base and added to PBS for further dissection. Next, the skin was removed, the femoral heads displaced from the acetabulum and the entire hind limbs detached and added to PBS. The skin was then removed from the tail and the tail tendons dissected free. Excess muscle was removed from the hind limbs and the feet removed at the tarsus. For techniques requiring isolated tendon and bone tissue, the patellar tendon was further removed, the femur and tibia separated, and all muscle dissected out.

### Pulse-chase with ^14^C-L-proline

Tail tendon, patellar tendon and femur tissue was dissected from 8 week old Tm1b mice. Tissue was dissected into small pieces and added to DMEM containing penicillin/streptomycin (1% v/v), L-glutamine (2 mM), L-ascorbic acid 2-phosphate (200 μM), β-aminopropionitrile (400 μM) and 2.5 μCi/ml [^14^C]proline (GE Healthcare, Illinois, USA) and incubated at 37°C for 18 hours. The tissue samples were subsequently moved to media without [^14^C]proline for 3 hours. Collagen was then extracted from the tissue samples using a salt extraction buffer (1 M NaCl, 25 mM EDTA, 50 mM Tris-HCl, pH 7.4) containing protease inhibitors (Roche, Basel, Switzerland). Samples were extracted overnight at 4°C with agitation. Extracts were analysed by electrophoresis on 6% Tris-Glycine gels (ThermoFisher, Massachusetts, USA) with delayed reduction (60). The gels were fixed (10% methanol, 10% acetic acid), dried under vacuum, and exposed to a phosphorimaging plate (BAS-IP MS). Phosphorimaging plates were processed using a phosphorimager (Typhoon FLA7000 IP) and densitometry carried out using ImageQuant software (GE Healthcare Life Sciences, Illinois, USA).

### Three point bending

Before three-point bending tests, the freshly isolated intact femurs and tibias (only those showing no evidence of fracture calluses) were imaged using uCT in order to obtain the cross-sectional area, circumference and moment of inertia measurements. Bones were scanned inside 1 ml syringes in PBS using a Skyscan 1272 scanner (Bruker, Kontich, Belgium). Scans were performed at a resolution of 9 μm (60 kV, 150 μA, 2×2 binning, rotation step size 0.5°, using a 0.5 mm aluminium filter). Scans were reconstructed using NRecon (Bruker, Kontich, Belgium) using Gaussian smoothing of 1, ring artefact reduction 5, and beam hardening compensation at 38%. For analysis of cortical bone parameters, a region of interest of 200 slices of the mid-femur and mid-tibia was selected and saved using Dataviewer (Bruker, Kontich, Belgium). Cross-sectional area for both bones, then cortical thickness, bone perimeter, periosteal circumference, second moment of area about the mediolateral axis and polar moment of inertia as well as bone density measurements for the femur, were obtained using custom macros in CTan (Bruker, Kontich, Belgium). Endosteal circumference was calculated using equation 1.1. Density measurements were calibrated using a set of hydroxyapatite phantoms (Bruker, Kontich, Belgium).

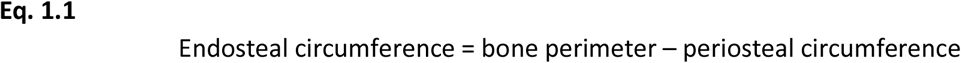

A Zwickiline Fmax 1 kN (Zwick, Ulm, Germany) biomechanical tester fitted with a 50N load cell was used for three-point bending experiments. Femurs and tibias were loaded at a span length of 8 mm and the crosshead was lowered at a rate of 0.5 mm/min using testXpert II software (Zwick). Ultimate force and stiffness measurements were calculated from the force-displacement curve, at the point of maximum load and the maximum gradient of the linear rising section of the graph respectively. The maximum stiffness and elastic modulus. were calculated as described (61)

### Micro computed tomography (uCT)

Hind limbs were fixed overnight in 10% neutral buffered formalin before washing and storage in 70% ethanol. Limbs were loaded into 2 ml syringe tubes in 70% ethanol and scanned using a Skyscan 1272 scanner (Bruker, Kontich, Belgium) at a resolution of 4.5 μm(60 kV, 150 μA, no binning, rotation step size 0.3°, using a 0.5 mm aluminium filter). Scans were reconstructed using NRecon software (Bruker) as described above. Trabecular bone parameters of the proximal tibia were measured in a volume, selected and saved using Dataviewer (Bruker), of 200 slices starting 20 slices distal to the growth plate as described (62). As previously, only bones that did not contain any fracture calluses were used. Trabecular bone parameters were measured using a custom macro in CTan (Bruker).

### Allelic discrimination

RNA was extracted from tissues preserved in RNAlater by firstly applying Trizol to samples which were then homogenised using a steel ball lysing matrix and a FastPrep 24 tissue homogeniser (MP Biomedicals, California, USA). RNA was extracted from homogenised samples using a Direct-Zol RNA kit (Zymo Research, California, USA) as per the manufacturer’s instructions. The quantity and quality of RNA was assessed using a NanoDrop spectrophotometer (Thermo Fisher, Massachusetts, USA), 260/280 values between 1.8-2.1 were deemed of a sufficient RNA quality. cDNA was synthesised in a 25 μl reaction from 0.5-1 μg of total RNA. The conditions for cDNA synthesis were: incubation at 5 minutes at 70°C, 60 minutes at 37°C and 5 minutes at 93°C with 1 U/μl RNasin ribonuclease inhibitor, 2 mM PCR nucleotide mix, 8 U/μl M-MLV reverse transcriptase and 0.02 μg/μl random-hexamer oligonucleotides per reaction.

Detection of murine Col1a2 wild-type and mutant alleles was performed using a custom snpsig™ real-time PCR mutation detection/allelic discrimination kit (Primerdesign, Southampton, UK). 10 ng of cDNA was added to 10 μl of PrecisionPLUS mastermix (Primerdesign, Southampton, UK), 1 μl of the custom genotyping primer/probe mix and 4 μl nuclease free water per reaction. Amplification was performed on a Stratagene qPCR machine with an initial enzyme activation step of 2 minutes at 95°C followed by 10 cycles of denaturation for 10 seconds at 95°C and extension for 60 seconds at 60°C. Finally, 35 cycles of denaturation for 10 seconds at 95°C and extension for 60 seconds at 68°C, with fluorogenic data collected during this extension step for the ROX (wild type) and VIC (oim) channels.

### Statistical analysis

Sample size calculations were carried out using G*Power 3.1.9.2 and Stata13 to give a power of at least 90% at the 5% level of significance. Primary outcomes were defined as bone stiffness (N/mm), bone volume (%) and trabecular separation (μm) with a standardised effect size of 2 deemed to be biologically important. For comparison, effect sizes were calculated from previously reported oim data (48) as −2.0 (41%) for femur stiffness, −2.8 (48%) for bone volume and 2.8 (66%) for trabecular separation. Comparison of ‘bone mineral density’ (DEXA) effect sizes for the oim (63) and Col1a2 null lines (53) at 14 wks indicated effect sizes of a similar magnitude (d=-1.6 and 1.7 respectively). Group sizes of 3 were calculated for two-way ANOVA on normally distributed data to test the effect of genotype (Stata13). The sample size in each group was increased by 20% to allow for non-normality of the data to give planned group sizes of 4.

All statistical analysis was completed using SigmaPlot 14.0 or 14.5 software. Comparisons of continuous measurements across sex and genotype were carried out using a two-way ANOVA with a Holm-Sidak post-hoc test. Where residuals did not meet assumptions of normality (Shapiro-Wilk test) or equal variance (Brown-Forsythe test), data were transformed with a Box-Cox or Johnson transformation using Mintab 18 before analysis. If a suitable transformation was not identified, male and female datasets were analysed separately using a one-way ANOVA (denoted with coloured font on graphs). If residuals still did not meet assumptions a nonparametric Kruskal-Wallis one-way ANOVA on ranks with a Dunn’s post-hoc test was used for comparisons (italicised on graphs). Comparisons of two categorical variables were done via a chi-squared test with the expected counts in each cell of the table being at least 5. The time to deterioration data were summarised via survival curves and statistically compared via a log-rank (Mantel-Cox) test, overall p-value is reported. No data points were excluded from statistical analysis. Data were plotted using GraphPad Prism 8.

## Author Contributions

KJL: Data curation, Formal Analysis, Investigation, Project administration, Supervision, Visualization, Writing – original draft, Writing – review & editing; LR: Data curation, Investigation, Visualization; GBG: Funding acquisition, Methodology, Writing – review & editing; PC: Conceptualization, Funding acquisition, Writing – review & editing; RA: Funding acquisition, Methodology, Writing – review & editing; GC: Formal Analysis, Funding acquisition, Writing – review & editing; RVH: Data curation, Investigation, Funding acquisition, Investigation, Methodology, Software, Supervision, Visualization, Writing – review & editing; EGC-L: Conceptualization, Formal Analysis, Funding acquisition, Project administration, Supervision, Visualization, Writing – original draft, Writing – review & editing.

## Acknowledgements

The study was funded by the UK Medical Research Council (MR/R00319X/1). LR was supported by the Erasmus+ program.

## Supplementary Data

**Figure S1.**
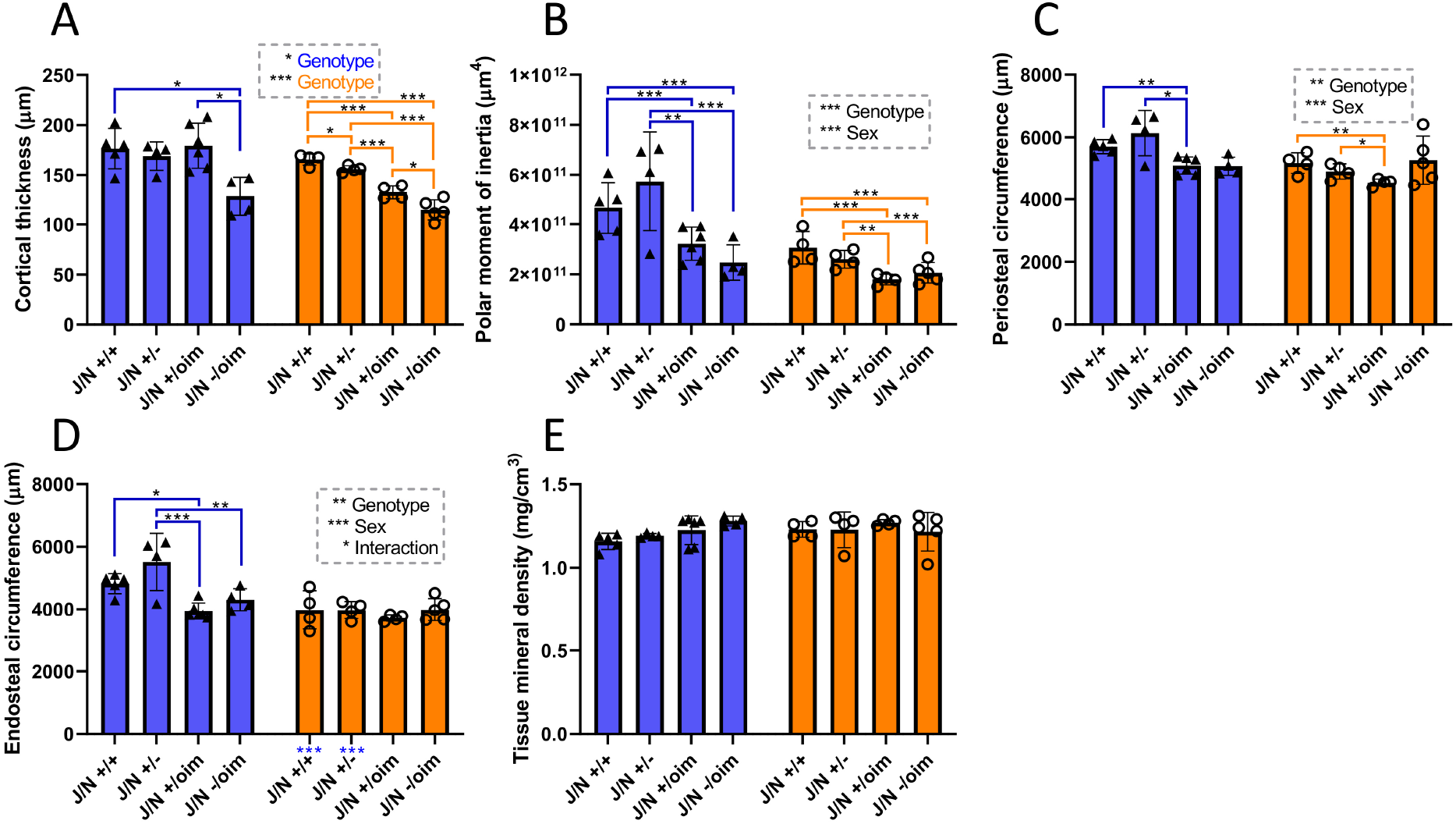
Femoral cortical bone analyses in mixed heterozygotes. MicroCT scans were performed at 8 weeks of age in the crossed oim/Col1a2 null line. Reconstruction and analysis of scan files enabled determination of cortical thickness (A), polar moment of inertia (B), periosteal (C) and endosteal (D) circumference, as well as bone density (E). Blue bars/triangles = males, orange bars/circles = females. Blue and orange brackets show differences between genotypes within males (blue) or females (orange). Male/female differences for particular genotypes are shown below the x-axis for females (blue stars). * p-value < 0.05, ** p-value <0.01 and *** p-value <0.001. n=4 for all groups, except male J/N +/+ and female J/N -/oim where n=5, and male J/N +/oim where n=6.

**Figure S2.**
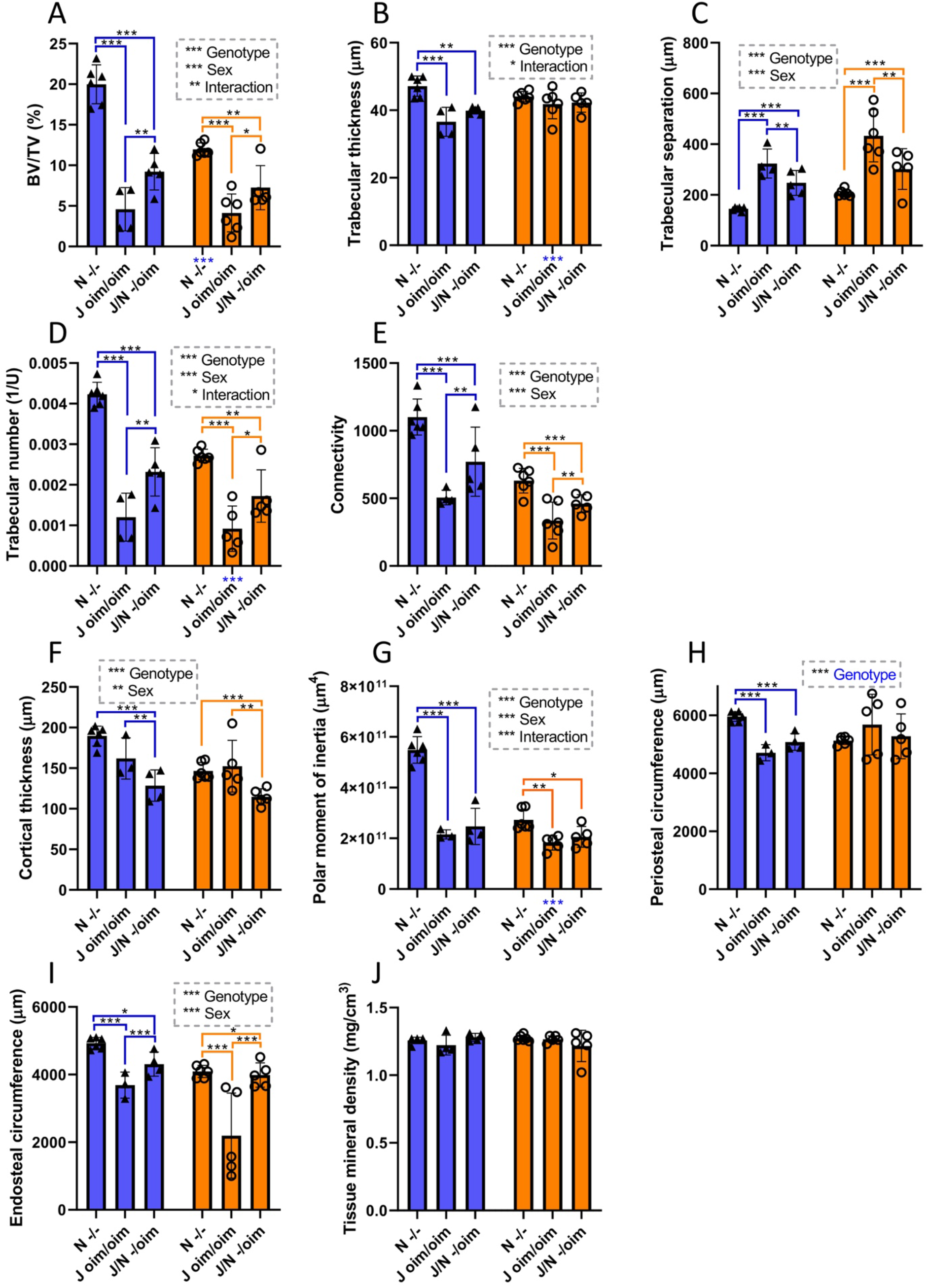
Bone structural properties in compound heterozygotes are generally less severe than in oim homozygotes. Bone volume (A), trabecular thickness (B), trabecular separation (C), trabecular number (D), connectivity (E), cortical thickness (F), polar moment of inertia (G), periosteal (H) and endosteal (I) circumference, as well as bone density (J) comparisons are shown for Col1a2 null homozygotes (-/-), oim homozygotes (oim/oim) and compound heterozygotes (-/oim). Blue bars/triangles = males, orange bars/circles = females. Blue and orange brackets show differences between genotypes within males (blue) or females (orange). Male/female differences for particular genotypes are shown below the x-axis for females (blue stars). * p-value < 0.05, ** p-value <0.01 and *** p-value <0.001. For trabecular parameters (A-E) n=6 for all groups except -/oim (both sexes) and female oim/oim in D where n=5, and for male oim/oim where n=4. For cortical parameters (F-J) n=6 for -/-, n=5 for female oim/oim and -/oim, whilst n=4 for male -/oim and n=3 for male oim/oim (except in J where n=4 for male oim/oim).

## References

1. Canty EG, Kadler KE. Procollagen trafficking, processing and fibrillogenesis. J Cell Sci. 2005;118(Pt 7):1341–53.

2. Lees JF, Tasab M, Bulleid NJ. Identification of the molecular recognition sequence which determines the type-specific assembly of procollagen. EMBO J. 1997;16(5):908–16.

3. Sharma U, Carrique L, Vadon-Le Goff S, Mariano N, Georges RN, Delolme F, et al. Structural basis of homo- and heterotrimerization of collagen I. Nat Commun. 2017;8:14671.

4. DiChiara AS, Li RC, Suen PH, Hosseini AS, Taylor RJ, Weickhardt AF, et al. A cysteine-based molecular code informs collagen C-propeptide assembly. Nat Commun. 2018;9(1):4206.

5. Ralston SH, Uitterlinden AG, Brandi ML, Balcells S, Langdahl BL, Lips P, et al. Large-scale evidence for the effect of the COLIA1 Sp1 polymorphism on osteoporosis outcomes: the GENOMOS study. PLoS Med. 2006;3(4):e90.

6. Philp AM, Collier RL, Grover LM, Davis ET, Jones SW. Resistin promotes the abnormal Type I collagen phenotype of subchondral bone in obese patients with end stage hip osteoarthritis. Scientific reports. 2017;7(1):4042.

7. Kerns JG, Gikas PD, Buckley K, Shepperd A, Birch HL, McCarthy I, et al. Evidence from Raman Spectroscopy of a Putative Link Between Inherent Bone Matrix Chemistry and Degenerative Joint Disease. Arthritis & Rheumatology. 2014;66(5):1237–46.

8. Bailey AJ, Sims TJ, Knott L. Phenotypic expression of osteoblast collagen in osteoarthritic bone: production of type I homotrimer. Int J Biochem Cell Biol. 2002;34(2):176–82.

9. Zhong B, Huang D, Ma K, Deng X, Shi D, Wu F, et al. Association of COL1A1 rs1800012 polymorphism with musculoskeletal degenerative diseases: a meta-analysis. Oncotarget. 2017;8(43):75488–99.

10. Brull DJ, Murray LJ, Boreham CA, Ralston SH, Montgomery HE, Gallagher AM, et al. Effect of a COL1A1 Sp1 binding site polymorphism on arterial pulse wave velocity: an index of compliance. Hypertension. 2001;38(3):444–8.

11. Makareeva E, Han S, Vera JC, Sackett DL, Holmbeck K, Phillips CL, et al. Carcinomas Contain a Matrix Metalloproteinase–Resistant Isoform of Type I Collagen Exerting Selective Support to Invasion. Cancer Res. 2010;70(11):4366–74.

12. Rojkind M, Giambrone MA, Biempica L. Collagen Types in Normal and Cirrhotic Liver. Gastroenterol. 1979;76(4):710–9.

13. Ehrlich HP, Brown H, White BS. Evidence for type V and I trimer collagens in Dupuytren’s Contracture palmar fascia. Biochem Med. 1982;28(3):273–84.

14. Han S, Makareeva E, Kuznetsova NV, DeRidder AM, Sutter MB, Losert W, et al. Molecular mechanism of type I collagen homotrimer resistance to mammalian collagenases. J Biol Chem. 2010;285(29):22276–81.

15. Sims TJ, Miles CA, Bailey AJ, Camacho NP. Properties of collagen in OIM mouse tissues. Connect Tissue Res. 2003;44 Suppl 1:202–5.

16. Pfeiffer BJ, Franklin CL, Hsieh FH, Bank RA, Phillips CL. Alpha 2(I) collagen deficient oim mice have altered biomechanical integrity, collagen content, and collagen crosslinking of their thoracic aorta. Matrix Biol. 2005;24(7):451–8.

17. Carriero A, Zimmermann EA, Paluszny A, Tang SY, Bale H, Busse B, et al. How tough is brittle bone? Investigating osteogenesis imperfecta in mouse bone. J Bone Miner Res. 2014;29(6):1392–401.

18. Chipman SD, Sweet HO, McBride DJ, Jr., Davisson MT, Marks SC, Jr., Shuldiner AR, et al. Defective pro alpha 2(I) collagen synthesis in a recessive mutation in mice: a model of human osteogenesis imperfecta. PNAS. 1993;90(5):1701–5.

19. Carleton SM, McBride DJ, Carson WL, Huntington CE, Twenter KL, Rolwes KM, et al. Role of genetic background in determining phenotypic severity throughout postnatal development and at peak bone mass in Col1a2 deficient mice (oim). Bone. 2008;42(4):681–94.

20. Grabner B, Landis WJ, Roschger P, Rinnerthaler S, Peterlik H, Klaushofer K, et al. Age- and genotype-dependence of bone material properties in the osteogenesis imperfecta murine model (oim). Bone. 2001;29(5):453–7.

21. Vouyouka AG, Pfeiffer BJ, Liem TK, Taylor TA, Mudaliar J, Phillips CL. The role of type I collagen in aortic wall strength with a homotrimeric. J Vasc Surg. 2001;33(6):1263–70.

22. Misof K, Landis WJ, Klaushofer K, Fratzl P. Collagen from the osteogenesis imperfecta mouse model (oim) shows reduced resistance against tensile stress. J Clin Invest. 1997;100(1):40–5.

23. Phillips CL, Pfeiffer BJ, Luger AM, Franklin CL. Novel collagen glomerulopathy in a homotrimeric type I collagen mouse (oim). Kidney Int. 2002;62(2):383–91.

24. Nicholls AC, Pope FM, Schloon H. Biochemical heterogeneity of osteogenesis imperfecta: New variant. Lancet. 1979;1(8127):1193.

25. Nicholls AC, Osse G, Schloon HG, Lenard HG, Deak S, Myers JC, et al. The clinical features of homozygous alpha 2(I) collagen deficient osteogenesis imperfecta. J Med Genet. 1984;21(4):257–62.

26. Pace JM, Wiese M, Drenguis AS, Kuznetsova N, Leikin S, Schwarze U, et al. Defective C-propeptides of the proalpha2(I) chain of type I procollagen impede molecular assembly and result in osteogenesis imperfecta. J Biol Chem. 2008;283(23):16061–7.

27. Saban J, Zussman MA, Havey R, Patwardhan AG, Schneider GB, King D. Heterozygous oim mice exhibit a mild form of osteogenesis imperfecta. Bone. 1996;19(6):575–9.

28. Camacho NP, Hou L, Toledano TR, Ilg WA, Brayton CF, Raggio CL, et al. The material basis for reduced mechanical properties in oim mice bones. J Bone Miner Res. 1999;14(2):264–72.

29. Yao X, Carleton SM, Kettle AD, Melander J, Phillips CL, Wang Y. Gender-dependence of bone structure and properties in adult osteogenesis imperfecta murine model. Ann Biomed Eng. 2013;41(6):1139–49.

30. Prockop DJ. Osteogenesis imperfecta. A model for genetic causes of osteoporosis and perhaps several other common diseases of connective tissue. Arthritis Rheum. 1988;31(1):1–8.

31. Mann V, Hobson EE, Li B, Stewart TL, Grant SFA, Robins SP, et al. A COL1A1 Sp1 binding site polymorphism predisposes to osteoporotic fracture by affecting bone density and quality. J Clin Invest 2001;107(7):899–907.

32. Malfait F, Symoens S, Coucke P, Nunes L, De Almeida S, De Paepe A. Total absence of the alpha2(I) chain of collagen type I causes a rare form of Ehlers-Danlos syndrome with hypermobility and propensity to cardiac valvular problems. J Med Genet. 2006;43(7):e36.

33. Guarnieri V, Morlino S, Di Stolfo G, Mastroianno S, Mazza T, Castori M. Cardiac valvular Ehlers-Danlos syndrome is a well-defined condition due to recessive null variants in COL1A2. Am J Med Genet A. 2019;179(5):846–51.

34. Maruelli S, Besio R, Rousseau J, Garibaldi N, Amiaud J, Brulin B, et al. Osteoblasts mineralization and collagen matrix are conserved upon specific Col1a2 silencing. Matrix Biol Plus. 2020;6–7:100028.

35. Makareeva E, Aviles NA, Leikin S. Chaperoning osteogenesis: new protein-folding disease paradigms. Trends Cell Biol. 2011;21(3):168–76.

36. Forlino A, Cabral WA, Barnes AM, Marini JC. New perspectives on osteogenesis imperfecta. Nat Rev Endocrinol. 2011;7(9):540–57.

37. Chessler SD, Byers PH. BiP binds type I procollagen pro alpha chains with mutations in the carboxyl-terminal propeptide synthesized by cells from patients with osteogenesis imperfecta. J Biol Chem. 1993;268(24):18226–33.

38. Lisse TS, Thiele F, Fuchs H, Hans W, Przemeck GK, Abe K, et al. ER stress-mediated apoptosis in a new mouse model of osteogenesis imperfecta. PLoS Genet. 2008;4(2):e7.

39. Forlino A, Kuznetsova NV, Marini JC, Leikin S. Selective retention and degradation of molecules with a single mutant alpha1(I) chain in the Brtl IV mouse model of OI. Matrix Biol. 2007;26(8):604–14.

40. Mirigian LS, Makareeva E, Mertz EL, Omari S, Roberts-Pilgrim AM, Oestreich AK, et al. Osteoblast Malfunction Caused by Cell Stress Response to Procollagen Misfolding in alpha2(I)-G610C Mouse Model of Osteogenesis Imperfecta. J Bone Miner Res. 2016;31(8):1608–16.

41. Gioia R, Tonelli F, Ceppi I, Biggiogera M, Leikin S, Fisher S, et al. The chaperone activity of 4PBA ameliorates the skeletal phenotype of Chihuahua, a zebrafish model for dominant osteogenesis imperfecta. Hum Mol Genet. 2017;26(15):2897–911.

42. Kimura S. Wide distribution of the skin type I collagen alpha 3 chain in bony fish. Comp Biochem Physiol B. 1992;102(2):255–60.

43. Exposito JY, Valcourt U, Cluzel C, Lethias C. The fibrillar collagen family. International journal of molecular sciences. 2010;11(2):407–26.

44. McBride DJ, Jr., Choe V, Shapiro JR, Brodsky B. Altered collagen structure in mouse tail tendon lacking the alpha 2(I) chain. J Mol Biol. 1997;270(2):275–84.

45. Kuznetsova NV, McBride DJ, Leikin S. Changes in thermal stability and microunfolding pattern of collagen helix resulting from the loss of alpha2(I) chain in osteogenesis imperfecta murine. J Mol Biol. 2003;331(1):191–200.

46. Han S, McBride DJ, Losert W, Leikin S. Segregation of type I collagen homo- and heterotrimers in fibrils. J Mol Biol. 2008;383(1):122–32.

47. Ranzoni AM, Corcelli M, Hau KL, Kerns JG, Vanleene M, Shefelbine S, et al. Counteracting bone fragility with human amniotic mesenchymal stem cells. Scientific reports. 2016;6:39656.

48. Vanleene M, Saldanha Z, Cloyd KL, Jell G, Bou-Gharios G, Bassett JH, et al. Transplantation of human fetal blood stem cells in the osteogenesis imperfecta mouse leads to improvement in multiscale tissue properties. Blood. 2011;117(3):1053–60.

49. Vanleene M, Porter A, Guillot P-V, Boyde A, Oyen M, Shefelbine S. Ultra-structural defects cause low bone matrix stiffness despite high mineralization in osteogenesis imperfecta mice. Bone. 2012;50(6):1317–23.

50. Bart ZR, Hammond MA, Wallace JM. Multi-scale analysis of bone chemistry, morphology and mechanics in the oim model of osteogenesis imperfecta. Connect Tissue Res. 2014;55 Suppl 1:4–8.

51. Miller E, Delos D, Baldini T, Wright TM, Pleshko Camacho N. Abnormal mineral-matrix interactions are a significant contributor to fragility in oim/oim bone. Calcif Tissue Int. 2007;81(3):206–14.

52. Zimmerman SM, Heard-Lipsmeyer ME, Dimori M, Thostenson JD, Mannen EM, O’Brien CA, et al. Loss of RANKL in osteocytes dramatically increases cancellous bone mass in the osteogenesis imperfecta mouse (oim). Bone Rep. 2018;9:61–73.

53. IMPC. International Mouse Phenotyping Consortium: Col1a2 [Available from: https://www.mousephenotype.org/data/genes/MGI:88468.

54. Simon MM, Greenaway S, White JK, Fuchs H, Gailus-Durner V, Wells S, et al. A comparative phenotypic and genomic analysis of C57BL/6J and C57BL/6N mouse strains. Genome Biol. 2013;14(7):R82.

55. Takigawa S, Frondorf B, Liu S, Liu Y, Li B, Sudo A, et al. Salubrinal improves mechanical properties of the femur in osteogenesis imperfecta mice. J Pharmacol Sci. 2016;132(2):154–61.

56. Oganesian A, Au S, Horst JA, Holzhausen LC, Macy AJ, Pace JM, et al. The NH2-terminal propeptide of type I procollagen acts intracellularly to modulate cell function. J Biol Chem. 2006;281(50):38507–18.

57. Hayata T, Nakamoto T, Ezura Y, Noda M. Ciz, a transcription factor with a nucleocytoplasmic shuttling activity, interacts with C-propeptides of type I collagen. Biochem Biophys Res Commun. 2008;368(2):205–10.

58. Marongiu M, Deiana M, Marcia L, Sbardellati A, Asunis I, Meloni A, et al. Novel action of FOXL2 as mediator of Col1a2 gene autoregulation. Dev Biol. 2016;416(1):200–11.

59. Skarnes WC, Rosen B, West AP, Koutsourakis M, Bushell W, Iyer V, et al. A conditional knockout resource for the genome-wide study of mouse gene function. Nature. 2011;474(7351):337–42.

60. Sykes B, Puddle B, Francis M, Smith R. The estimation of two collagens from human dermis by interrupted gel electrophoresis. Biochem Biophys Res Commun. 1976;72(4):1472–80.

61. Schriefer JL, Robling AG, Warden SJ, Fournier AJ, Mason JJ, Turner CH. A comparison of mechanical properties derived from multiple skeletal sites in mice. Journal of biomechanics. 2005;38(3):467–75.

62. van ‘t Hof RJ, Dall’Ara E. Analysis of Bone Architecture in Rodents Using Micro-Computed Tomography. Methods Mol Biol. 2019;1914:507–31.

63. Phillips CL, Bradley DA, Schlotzhauer CL, Bergfeld M, Libreros-Minotta C, Gawenis LR, et al. Oim mice exhibit altered femur and incisor mineral composition and decreased bone mineral density. Bone. 2000;27(2):219–26.

